# Apolipoprotein E intersects with amyloid-β within neurons

**DOI:** 10.1101/2022.12.13.520238

**Authors:** Sabine C Konings, Emma Nyberg, Isak Martinsson, Laura Torres-Garcia, Claudia Guimas Almeida, Gunnar K Gouras

## Abstract

Apolipoprotein E4 (ApoE4) is the most important genetic risk factor for Alzheimer’s disease (AD). Among the earliest changes in AD is endosomal enlargement in neurons, which was reported as enhanced in ApoE4 carriers. ApoE is thought to be internalized into endosomes of neurons, while β-amyloid (Aβ) accumulates within neuronal endosomes early in AD. However, it remains unknown whether ApoE and Aβ intersect intracellularly. We show that internalized astrocytic ApoE localizes mostly to lysosomes in neuroblastoma cells and astrocytes, while in neurons it preferentially localizes to endosomes-autophagosomes of neurites. In AD transgenic neurons, astrocyte-derived ApoE intersects intracellularly with amyloid precursor protein (APP)/Aβ. Moreover, ApoE4 increases the levels of endogenous and internalized Aβ_42_ in neurons. Taken together, we demonstrate differential localization of ApoE in neurons, astrocytes and neuron-like cells, and show that internalized ApoE intersects with APP/Aβ in neurons, which may be of considerable relevance to AD.

## Introduction

The apolipoprotein E (ApoE) allele exists as three major isoforms in humans: ApoE2, ApoE3 and ApoE4, with ApoE3 being the most common form, followed by ApoE4. ApoE4 is the most important genetic risk factor for Alzheimer’s disease (AD)^1,2^. In contrast, ApoE2 protects against AD. ApoE in the brain is made predominantly by astrocytes, but is also generated by microglia and under conditions of stress by neurons^3,4^. Although the most critical mechanism(s) by which ApoE4 raises the risk of AD remains to be determined, various hypotheses have been proposed. A tight correlation of ApoE4 with amyloid pathology is well known, but the mechanism(s) whereby ApoE4 impacts amyloid pathology in AD remains unclear. Previous research suggests a role of ApoE4 in the early stages of AD, prior to amyloid plaques^5,6^. The presence of ApoE4 increases the number of dystrophic neurites around plaques and affects initial plaque density, but does not appear to influence plaque load after initial plaque formation^5,6^, suggesting a role of ApoE4 in the early cellular phase of AD prior to amyloid plaques.

Endosomal enlargement in neurons is one of the earliest changes associated with AD^7,8^. Post-mortem brains from late-onset AD patients showed significantly enlarged neuronal endosomes compared to age-matched control brains^7^, a finding that has also been described *in vitro* in IPSC-induced neurons from AD patients^9^ and as induced by AD genetic risk factors^10,11^. The mean endosomal size was reported as even more increased in post-mortem AD human brains from ApoE4 carriers compared to non-ApoE4 carriers, suggesting an enhancing effect of ApoE4 on endosomal enlargement^7^. The exact mechanism(s) underlying endosome dysfunction in AD, in particular in relation to ApoE4, remains poorly understood. Experimental studies have reported endosomal dysregulation induced by ApoE4 and ApoE4-induced impairment in endosome recycling^12–15^. Added recombinant ApoE was described to localize in neurons to the endosomal system at Rab5-positive early endosomes and cathepsin D-positive late endosomes/lysosomes^16^. However, studies on the endosomal localization of more physiological lipidated ApoE isoforms in neurons are lacking.

Prior to plaque formation, intraneuronal Aβ accumulates in endosomes^17–19^, in particularly in late endosomal multivesicular bodies (MVBs) near synaptic compartments^18^. In addition, BACE1-induced APP cleavage is favored under acidic pH conditions as present in endosomes^20^, APP trafficking and Aβ production occur in endosomes^11,21,22^. As Aβ is generated^21,23^ and initially aggregates^18^ in acidic endosomes, ApoE4 might facilitate the optimal conditions for Aβ production by blocking endosome recycling or provide a more favorable lipid environment for Aβ aggregation. ApoE was reported to trigger APP endocytosis and subsequent Aβ production after binding to ApoER2 receptors, with ApoE4 increasing Aβ production more than ApoE2 and ApoE3^24^. *In vivo* data also support an association of ApoE4 with increased levels of Aβ, as human ApoE target replacement mice transduced with lentiviral Aβ_1-42_ showed increased intracellular Aβ deposition in ApoE4 compared to ApoE3 mouse brains^25^. On the other hand, in other studies mouse brain knock-in with the different human ApoE isoforms did not reveal alterations in APP levels or processing^26^.

It is possible that internalized ApoE and APP and/or Aβ intersect and even interact within endosomes. ApoE localizes with high molecular weight Aβ oligomers in the TBS-soluble fraction of human AD brain^27^, and ApoE co-deposits with amyloid plaques^28^, suggesting that ApoE and Aβ also co-localize at later stages of AD, but whether ApoE and APP/Aβ are intersecting at an intracellular level, remains less clear. ApoE and antibody 4G8-positive Aβ/APP was reported to be present in the same cytoplasmic granules in postmortem AD brains^29^, although the antibody used did not distinguish Aβ from its abundant precursor APP. A subsequent study using Aβ_42_ C-terminal-specific antibodies noted localization of ApoE to AD-vulnerable neurons with marked intraneuronal Aβ_42_ accumulation in human brains with early AD pathology^30^. More recently, ApoE and Aβ were reported to be present in the same synapses of human AD brains^31,32^.

Despite ApoE4 being established as the most important genetic risk factor for AD and neurons selectively degenerating in the disease, the subcellular localization of ApoE and its potential relation to Aβ/APP biology in neurons has not been explored. The aim of this study was to examine the subcellular localization of ApoE3 and ApoE4. In addition, the localization of the ApoE isoforms with intracellular Aβ/APP was examined. We demonstrate that internalized ApoE predominantly traffics to lysosomes in primary astrocytes and N2a neuroblastoma cells, but not in primary neurons where it localizes to endosomes and autophagosomes of neurites. Furthermore, ApoE co-localized with Aβ/APP β-C-terminal fragments (APP-βCTFs), with ApoE4 treated-neurons showing more prominent intraneuronal Aβ.

## Results

### ApoE is internalized into the endosome-lysosome system of N2a cells

The endosome-lysosome system is considered to be one of the earliest cellular sites affected in AD^7,33,34^. ApoE4 was previously shown to impact neuronal endosomes, including endosomal recycling^12^ and endosomal morphology^14^, however the subcellular biology of ApoE remains poorly studied. In order to better define the subcellular localization of ApoE, N2a neuroblastoma cells were initially treated with physiological levels (2.5 μg/ml) of recombinant human ApoE3 or ApoE4 for 15 min and 4 h (**Figure 1A**). After 4 h, recombinant ApoE3 and ApoE4 were internalized by the majority of N2a cells (**Figure 1B, Supplementary Figure 1A**) and were seen in a vesicle-like pattern (**Figure 1C**), consistent with the endosomal-lysosomal system, although interestingly, most of the ApoE labeling appeared polarized and near to the Golgi apparatus and microtubule organizing center (MTOC). Due to endogenous mouse ApoE expression by N2a cells and the fact that the ApoE antibody 16H22L18 to some extent also detects mouse ApoE, we were unable to distinguish differences in ApoE in vehicle-treated N2a cells from human ApoE-treated cells at the shorter (15 min) time point (data not shown). For this reason, further characterization of human ApoE in neuronal cells was performed using the 4 h time point, at which time the internalized ApoE was clearly above the lower endogenous mouse ApoE labeling.

**Figure 1:**
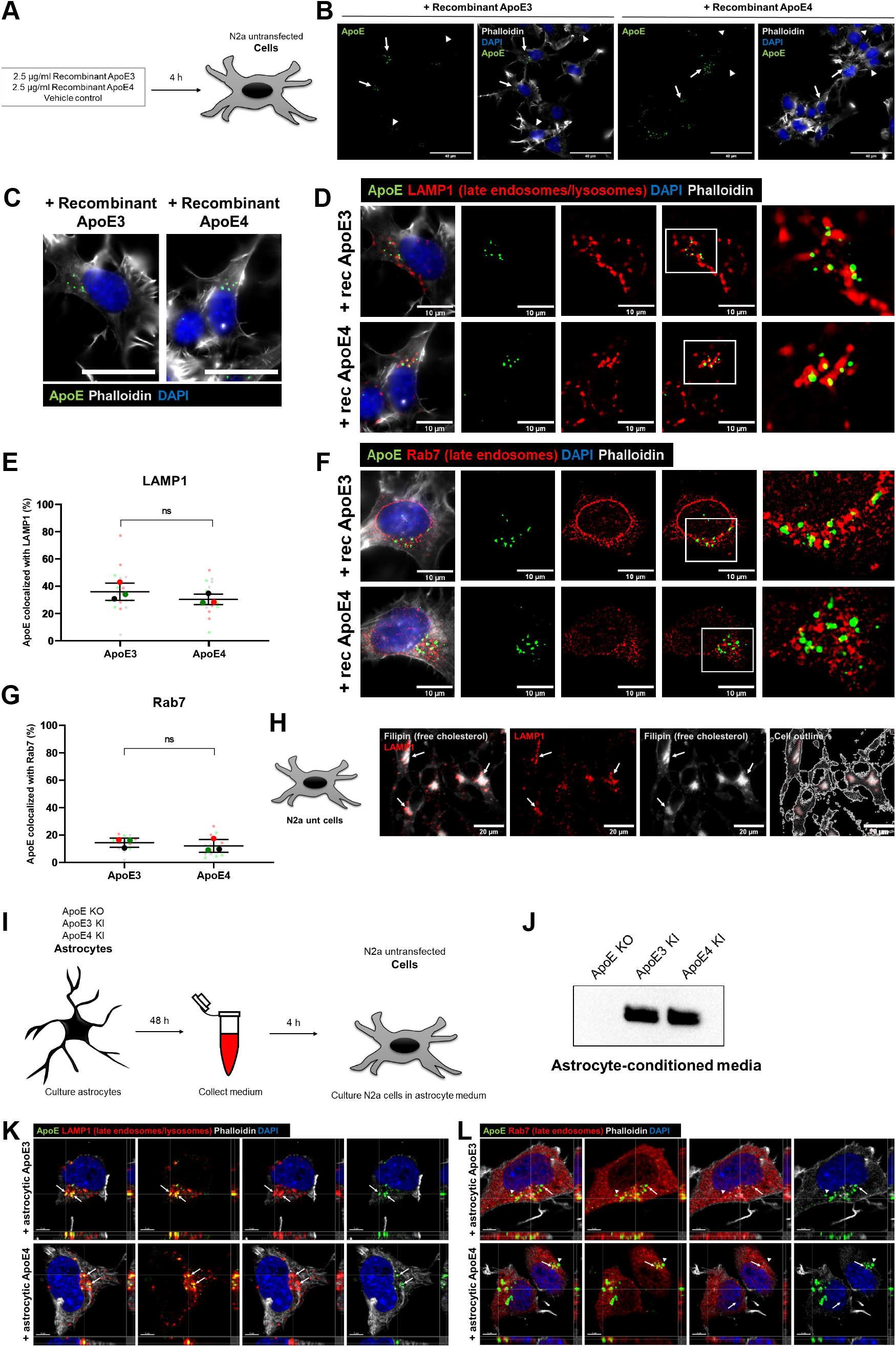
Recombinant and astrocyte-derived ApoE are internalized into the endosome-lysosome system in N2a cells. **A.** Schematic overview of 4 h treatment with 2.5 μg/ml recombinant ApoE3, ApoE4 or vehicle control in N2a cells. **B.** Representative epifluorescence images of N2a cells treated with recombinant ApoE3 or ApoE4 for 4 h showing an overview of human ApoE internalization in N2a cells. The N2a cells were labeled for ApoE (green), DAPI (blue) and phalloidin (grey). Cells showing internalized human ApoE were indicated by arrows, cells negative for ApoE were indicated by arrowheads. Scale bar is 40 μm. **C.** Higher magnification image of ApoE3- and ApoE4-treated N2a cells from **Figure 1B**. Internalized recombinant ApoE3 and ApoE4 puncta detected in N2a cells show a vesicle-like pattern. Scale bar represents 40 μm. **D.** Representative images obtained by epifluorescence microscopy of N2a cells incubated with recombinant ApoE3 or ApoE4 for 4 h. The cells were labeled for ApoE (green), late endosomal/lysosomal marker LAMP1 (red), DAPI (blue) and phalloidin (grey). The scale bar represents 10 μm. **E.** Quantification of **Figure 1D**, showing the co-localization levels (in percentages) of ApoE with LAMP1. **F.** Representative epifluorescence images of recombinant ApoE3- and ApoE4-treated N2a cells labeled for ApoE (green), late endosomal marker Rab7 (red), DAPI (blue) and phalloidin (grey). Scale bar is equal to 10 μm. **G.** Quantification of ApoE-Rab7 co-localization of **Figure 1F**. **H.** Representative images taken by epifluorescence microscopy of N2a cells showing LAMP1 (red) overlaps with Filipin (grey), a free cholesterol dye. Scale bar is 20 μm. **I.** Schematic representation of astrocyte-conditioned media collection from ApoE KO, ApoE3 KI and ApoE4 KI primary astrocytes, followed by media incubation of N2a cells for 4 h. **J.** Representative western blot of secreted human ApoE proteins detected in astrocyte conditioned media after culturing 48 h with ApoE KO, ApoE3 KI and ApoE4 KI primary astrocytes. **K-L.** Representative orthogonal images obtained by confocal microscopy showing ApoE (green) co-localizing with LAMP1 (red) **(K)** and to a lower extent also with Rab7 (red) **(L)** in ApoE3 and ApoE4 astrocyte conditioned media treated N2a cells. The cells were further labeled for DAPI (blue) and phalloidin (grey). Arrows indicate ApoE puncta inside LAMP1- **(K)** and Rab7- **(L)** positive vesicles. Scale bars represent 5 μm. Data is expressed as mean ± SD. ns = non-significant, Rec ApoE = recombinant ApoE. See also Supplementary Figure 1A-C.

To better define the presumed endosomal-lysosomal trafficking of internalized recombinant ApoE in N2a cells, we co-labeled ApoE-treated N2a cells for ApoE and select endosome-lysosome markers. Due to the vesicle-like localization of ApoE, we started with analyzing ApoE co-localization with antibodies to LAMP1, a lysosomal marker, which also detects late endosomes, and to Rab7, a late endosomal marker. At 4 h almost 40% of internalized human ApoE co-localized with LAMP1 (35-40% of ApoE-positive pixels overlap with LAMP1-positive pixels; **Figure 1D-E, Supplementary Figure 1B**), suggesting lysosomal and/or late endosomal localization of internalized recombinant ApoE3 and ApoE4 in N2a cells. No significant difference in ApoE and LAMP1 co-localization was detected between ApoE3 and ApoE4 (**Figure 1E**), implying no ApoE isoform-dependent differences in subcellular localization at 4 h. A lower proportion of human ApoE puncta co-localized with Rab7 in N2a cells after 4 h (around 15%; **Figure 1F-G**). We note that Rab7 labeling distributed more widely throughout the cell while most ApoE and LAMP1 localized in the perinuclear region, likely near the Golgi/MTOC. Overall, the LAMP1-positive, mostly Rab7-negative, co-labeling of internalized human ApoE suggests that human ApoE after 4 h has mostly trafficked beyond late endosomes to lysosomes in N2a cells. To assess whether internalized ApoE might also be transported to the Golgi apparatus, the presence of ApoE at the cis- and trans-Golgi apparatus was assessed. Despite preferential labeling of internalized ApoE3 and ApoE4 near to GM130-labeled cis-Golgi apparatus (**Supplementary Figure 2A**), no clear direct co-localization was seen of human ApoE with the antibody against GM130 nor with the trans-Golgi marker TGN38 (**Supplementary Figure 2**). Altogether internalized recombinant ApoE3 and ApoE4 in N2a cells are most notably localized to lysosomes after 4 h incubation.

Although recombinant ApoE is generally considered to be poorly lipidated, previous research suggests that cell media containing fetal bovine serum (FBS) can provide ApoE with lipids^35^, and we note that our N2a cell media contains FBS. Since ApoE is the main lipid and cholesterol carrier in the brain, we next examined whether cholesterol localizes to similar subcellular compartments as ApoE. Interestingly, filipin, a dye staining free cholesterol, revealed that in N2a cells cholesterol, analogous to recombinant ApoE, localized to LAMP1-positive vesicles (**Figure 1H**).

Cellular trafficking of ApoE might however differ depending on the lipidation of ApoE. Therefore, we next studied the cellular localization of ApoE under more physiological conditions by using ApoE particles obtained from humanized ApoE3-knock-in (KI) or ApoE4-KI primary astrocytes^36^. Astrocytes are the main source of ApoE in the brain^37^ and astrocyte-derived ApoE is lipidated^38^ and transported to other cell types including neurons. Human ApoE3 and ApoE4 astrocyte conditioned media were collected from ApoE3- and ApoE4-KI mouse primary astrocytes and subsequently used to treat N2a cells for 4 h (**Figure 1I**). Human ApoE was readily detected by Western blot in media collected from ApoE3- and ApoE4-KI mouse astrocytes, but, as expected, not from ApoE knock-out (KO) astrocytes (**Figure 1J**), confirming ApoE3 and ApoE4 conditioned media as a useful source for human ApoE. Like recombinant human ApoE, added astrocyte-derived human ApoE3 and ApoE4 after 4 h co-localized substantially with LAMP1-positive vesicles in N2a cells (**Figure 1K, Supplementary Figure 1C**), and showed lower levels of co-localization in the more numerous Rab7-positive late endosomes **(Figure 1L**). These findings confirm that internalized human ApoE3 and ApoE4, recombinant or derived from primary astrocytes, mostly traffics to lysosomes of N2a cells.

### Internalized ApoE in primary neurons preferentially localizes to endosomes and autophagosomes in neurites

Even though N2a cells are considered to be neuron-like cells, their morphology and cellular physiology are quite different from mature neurons. To study the localization of astrocyte-derived human ApoE in mature primary neurons, the subcellular localization of human ApoE was examined after adding conditioned astrocyte ApoE3 or ApoE4 media to ApoE KO primary brain cultures for 4 h (**Figure 2A**). ApoE KO brain cultures were used to allow only the visualization of the subcellular localization of added astrocyte-derived human ApoE without the presence (and background signal) of endogenous mouse ApoE. Since in N2a cells both recombinant and astrocyte-derived ApoE localized mostly to LAMP1-positive vesicles (**Figure 1**), we initially studied neuronal cell bodies, where lysosomes are predominantly located in neurons^39–41^. Surprisingly, after 4 h of treatment internalized astrocyte-derived ApoE3 and ApoE4 localized only to a limited extent to neuronal cell bodies where, unlike N2a cells, ApoE was not found to co-localize with LAMP1-positive vesicles (**Figure 2B**; ApoE KO media control in **Supplementary Figure 1D**), suggesting limited trafficking of added astrocytic ApoE to lysosomes in neuron soma. Internalized human ApoE puncta were also not seen to overlap with Rab7 in neuronal soma (**Figure 2C**). Of note, most ApoE labeling near to cell bodies actually appeared to be in the periphery and/or just outside the neuron cell soma.

**Figure 2:**
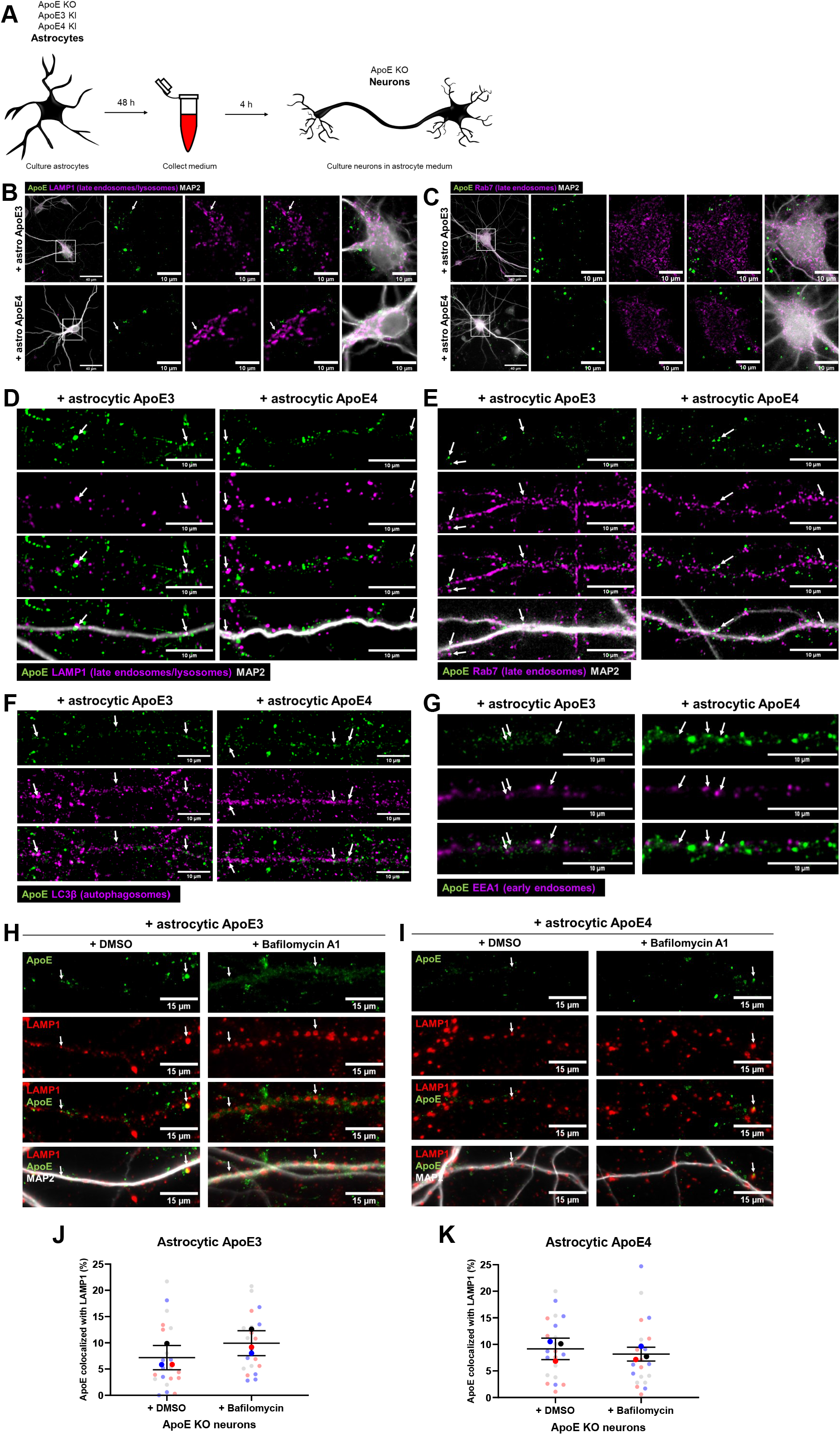
Internalized ApoE in primary neurons preferentially localizes to endosomes and autophagosomes in neurites. **A.** Schematic overview of ApoE KO, ApoE3 KI and ApoE4 KI astrocyte conditioned medium collection and subsequent 4 h treatment to ApoE KO primary neuron cultures (19 DIV). **B-C.** Representative fluorescent images obtained by epifluorescence microscopy of ApoE KO neurons treated with ApoE3 or ApoE4 astrocyte conditioned medium for 4 h with the focus on neuronal cell bodies. The ApoE astrocyte medium-treated neurons were labeled for ApoE (green), neuronal dendrite marker MAP2 (grey) and either LAMP1 **(B)** or Rab7 **(C)** (both shown in magenta). The left panels **(B-C)** show an overview of the entire neuron; higher magnification images are shown in the other four panels. The white arrows indicate ApoE puncta overlap with LAMP1-positive vesicles in neuronal cell bodies **(B)**. Left panels: scale bars are 40 μm. Right panels: scale bars are 10 μm. **D-G.** Representative epifluorescence images of neurites from ApoE3 and ApoE4 media-treated primary neurons. The neurites are labeled with ApoE (green), MAP2 **(D-E)** (grey), and late endosomal/lysosomal marker LAMP1 **(D)**, late endosomal marker Rab7 **(E)**, autophagosomal marker LC3β **(F)** or early endosomal marker EEA1 **(G)** (all in magenta). White arrows indicate co-localization between ApoE and the subcellular markers **(D-G)**. Scale bars represent 10 μm. **H-I.** Representative fluorescence images of ApoE KO neurites treated with control (DMSO) or 10 nM lysosomal inhibitor bafilomycin A1 for 1 h, followed by 4 h ApoE3 (**H**) or ApoE4 (**I**) astrocyte conditioned media. The neurites were labeled for ApoE (green), LAMP1 (red) and MAP2 (grey). ApoE and LAMP1 co-localization, indicating the presence of ApoE at late endosomes and/or lysosomes, is indicated by white arrows. Scale bar is 15 μm. **J-K.** Quantification of ApoE and LAMP1 co-localization in ApoE3-treated neurons shown in **Figure 2H** (**J**) and ApoE4-treated neurons shown in **Figure 2I** (**K**) with and without bafilomycin A1 treatment. The researcher performing the quantifications was blinded. Data is shown as mean ± SD. See also Supplementary Figure 1D-E.

Remarkably, in primary neurons (DIV 19) the majority of added astrocyte-derived ApoE after 4 h was seen in neurites rather than neuronal cell bodies (**Figure 2D-G**). In contrast to neuronal soma, added astrocytic ApoE3 and ApoE4 after 4 h were seen to be present in LAMP1-positive vesicles in neurites, which in our neurons should represent late endosomes based on their *in vitro* age (**Figure 2D; ApoE KO media control in Supplementary Figure 1E**)^42,43^. In line with this, the internalized human ApoE also partially colocalized with Rab7 in neurites **(Figure 2E**). However, the majority of added astrocytic ApoE did not colocalize with either LAMP1- or Rab7-positive puncta, highlighting other cellular site(s) of ApoE in neurites. Since internalized astrocytic ApoE3 and ApoE4 labeled in a vesicular pattern, the subcellular localization of ApoE in neurites was further studied using additional markers, with EEA1 to identify early endosomes, and LC3β to label autophagosomes. After 4 h added astrocytic ApoE was detected in both LC3β-positive autophagosomes (**Figure 2F**) and EEA1-positive early endosomes (**Figure 2G**) at comparable levels to LAMP1- and Rab7-positive vesicles (**Figure 2D-E**). Altogether, these findings suggest that astrocyte-derived ApoE, after internalization into primary neurons, is mostly present in the endosome-autophagy system of neurites.

Since levels of internalized ApoE in lysosomes of neuron soma were not as apparent as in N2a cells, we wondered whether this might also be caused by degradation of astrocyte-derived ApoE by neuronal lysosomes. To assess possible degradation of ApoE by lysosomes in neurons, we inhibited lysosomal degradation using 10 nM bafilomycin A1 (BafA1) starting 1 h prior to addition for 4 h of ApoE3 or ApoE4 conditioned media. BafA1 inhibits lysosomal function by blocking lysosomal acidification via v-ATPase^44,45^. The size of LAMP1-positive vesicles in neurites increased after addition of BafA1, suggesting that bafilomycin A1 was successfully blocking lysosomal function in our cultures (**Supplementary Figure 3**)^46^. If ApoE is degraded by neuronal lysosomes, it would be expected that the co-localization of ApoE with LAMP1 would increase after bafilomycin A1 treatment. However, no significant change in ApoE-LAMP1 co-localization was detected after BafA1 treatment for either ApoE3 or ApoE4 (**Figure 2H-K**), suggesting that lysosomal degradation in neurons is not a major pathway of internalized astrocyte-derived ApoE.

### Astrocyte-derived ApoE localizes to LAMP1-positive vesicles in primary astrocytes

While in MAP2-positive neurons some co-localization between internalized ApoE and LAMP1 was seen particularly in their neurites, we also observed that bright ApoE-positive puncta overlapped with LAMP1-positive vesicles in cells negative for the neuronal marker MAP2 (**Figure 3A**). Since MAP2 labels dendrites, antibody SMI-130 was used to assess whether this added human ApoE might be in axons. However, the strong ApoE-positive puncta did not clearly follow SMI-130 axonal labeling (**Supplementary Figure 4A**) and therefore appear not to be in axons.

**Figure 3:**
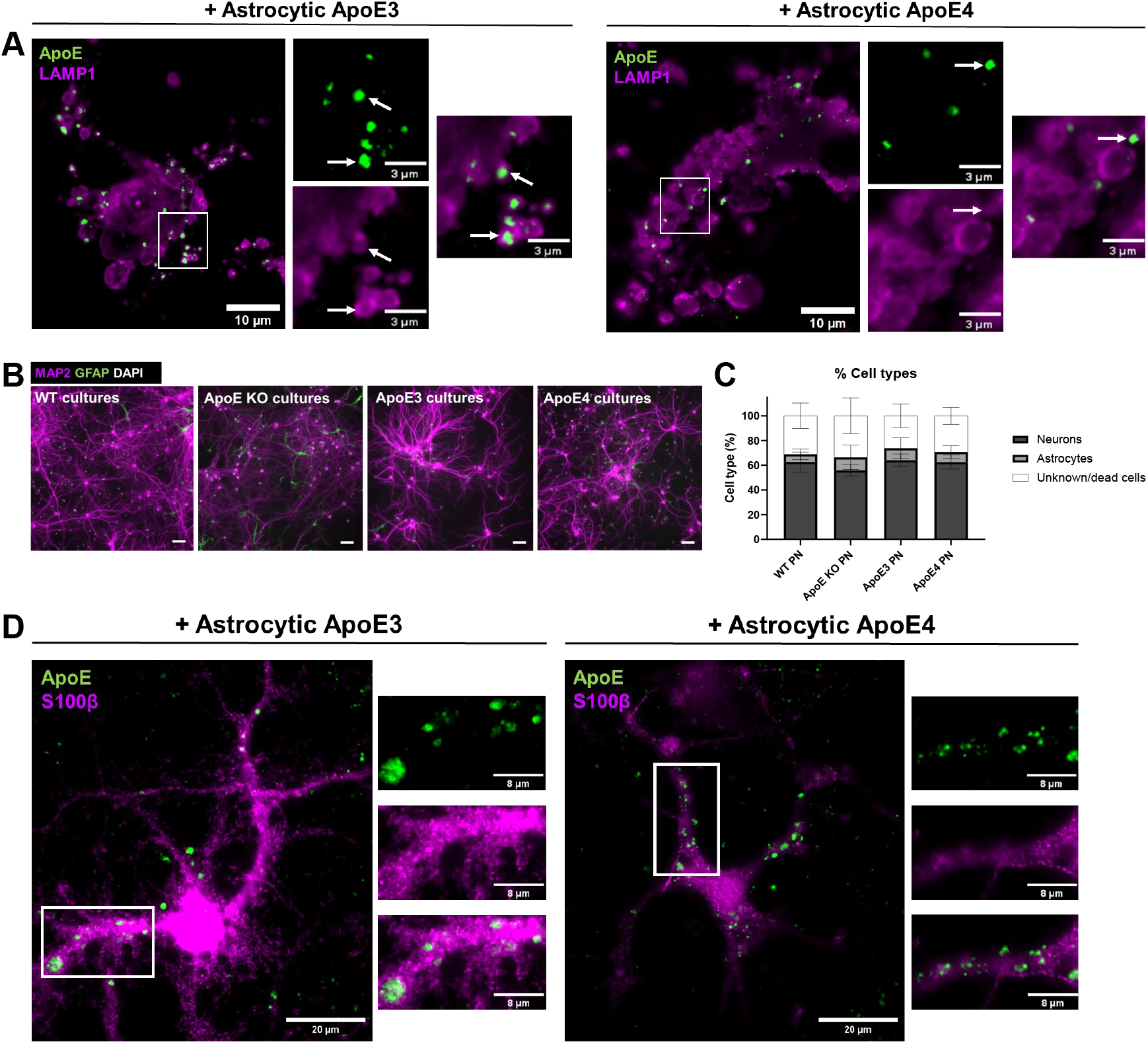
Astrocyte-derived ApoE localizes to LAMP1-positive vesicle in astrocytes. **A.** Representative fluorescence images of LAMP1-positive puncta (magenta) and ApoE (green) overlap in MAP2-negative cells in primary cultures. The primary brain cultures were treated with ApoE3 or ApoE4 astrocyte conditioned media for 4 h. The overlap of LAMP1 with human ApoE was highlighted by white arrows. Scale bars are 10 μm (bigger panels) and 3 μm (smaller panels). **B.** Representative images of wild-type, ApoE KO, ApoE3 KI and ApoE4 KI primary cultures labeled for neuronal marker MAP2 (magenta), astrocyte marker GFAP (green) and nuclear marker DAPI (white). Scale bar is 50 μm. **C.** Quantifications of MAP2-positive and GFAP-positive cells (%) in the primary cultures described in **Figure 3B**. The total number of cells was set based on the number of DAPI-positive nuclei present. Data is shown as mean ± SD. **D.** Representative fluorescence images of S100β-positive astrocytes (magenta) labeled for ApoE (green). The primary cultures containing the S100β-positive astrocytes were treated with ApoE3 or ApoE4 astrocyte conditioned media for 4 h. Scale bar represents 20 μm. WT = wild-type, PN = primary neurons.

To assess the presence of other cell types in our primary mouse brain cultures, which might relate to this strong extra-neuronal ApoE/LAMP1 labeling, the number of MAP2-postive neurons and GFAP-positive astrocytes were initially analyzed in relation to the DAPI nuclei present. Around 60% of all cells present in our cultures were positive for MAP2, indicative of neurons (**Figure 3B-C**). A minority of the cells in our cultures (approximately 5-10%) labeled positive for GFAP, suggesting a minority of the cell population represents astrocytes. However, 25-30% of all DAPI-nuclei were negative for MAP2 as well as GFAP. To further study other cell types, we labeled our wild-type primary brain cultures with antibodies to Iba1, to identify microglia, and CD140a, to label oligodendrocyte precursor cells (OPCs). As expected in embryonic brain cell cultures, no Iba1-positive microglia were detected (**Supplementary Figure 4B**). In contrast, there were numerous CD140a-positive OPCs (**Supplementary Figure 4C**). To study whether the ApoE is internalized by astrocytes or OPCs, ApoE KO mixed brain cultures were treated with ApoE3 and ApoE4 astrocyte conditioned media and co-labeled for human ApoE and astrocyte marker S100β or OPC marker CD140a. While added astrocytic ApoE was not clearly observed in CD140a-positive OPCs (**Supplementary Figure 4D**), added human ApoE clearly localized to S100β-positive astrocytes (**Figure 3D**) in a similar pattern as seen for LAMP1 (**Figure 3A**). Thus, primary astrocytes present in our ApoE KO mouse brain cultures readily take up added astrocyte-derived human ApoE where it is detected prominently in LAMP1 vesicles. Interestingly, similar to neurons, internalized ApoE puncta were mostly detected in astrocytic processes rather than cell soma (**Figure 4D**).

**Figure 4:**
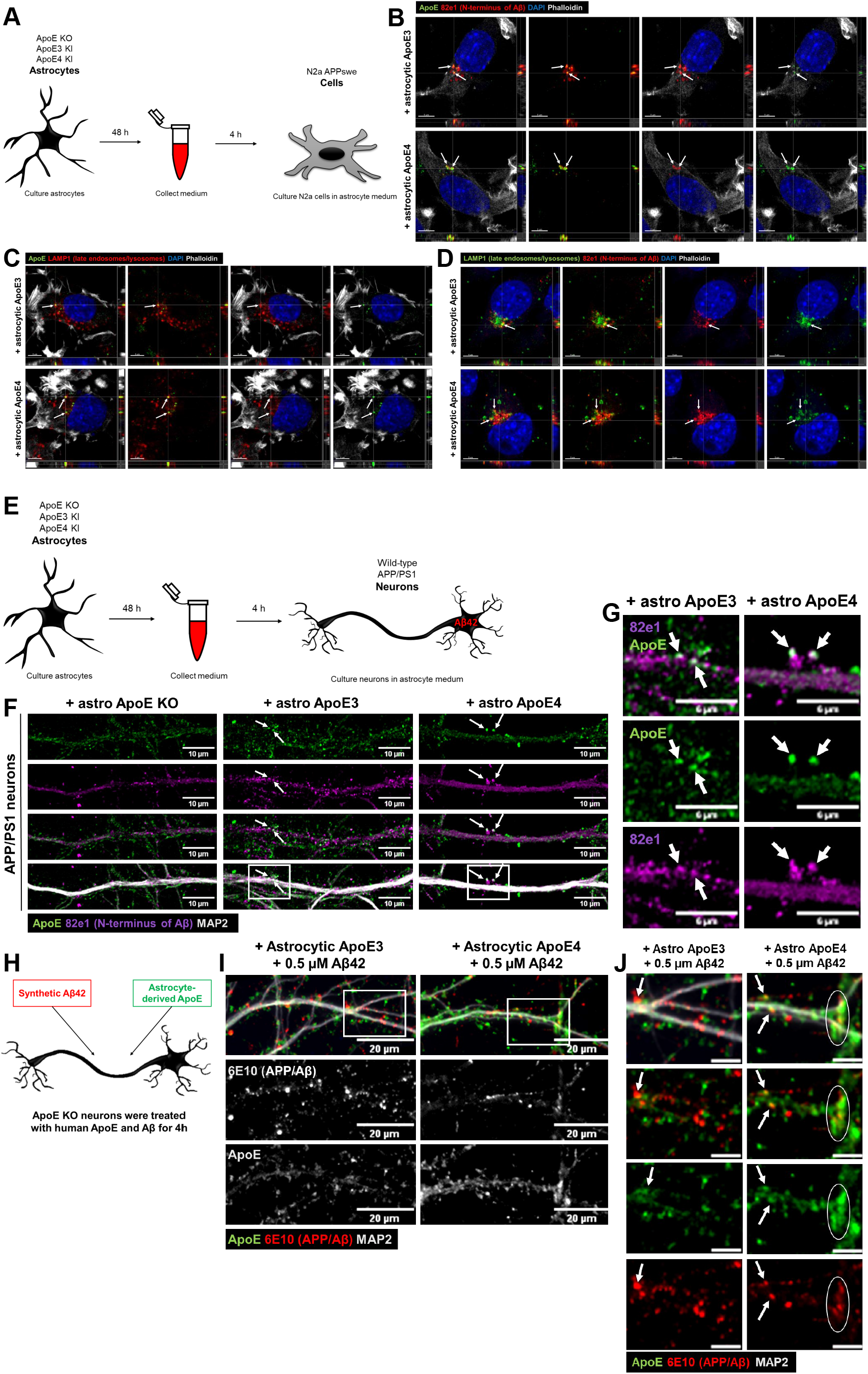
Internalized ApoE co-localizes with APP cleavage products in neurons and N2a cells. **A.** Schematic representation of experimental approach used. Astrocyte conditioned medium was collected from ApoE KO, ApoE3 KI and ApoE4 KI primary astrocytes and subsequently added to N2a APP_Swe_ cells for 4 h. **B.** Representative orthogonal images obtained by confocal microscopy of N2a APP_Swe_ cells incubated with ApoE3 and ApoE4 astrocyte conditioned media for 4 h. The cells were labeled for ApoE (green), human Aβ/APP-βCTFs by antibody 82e1 (red), DAPI (blue) and phalloidin (grey). Human ApoE and 82e1-positive Aβ/APP-βCTFs overlap as indicated by white arrows. **C.** Orthogonal fluorescent images of astrocytic ApoE3 and ApoE4-treated N2a APP_Swe_ cells. ApoE was labeled in green, late endosomes/lysosomes with LAMP1 in red, DAPI in blue and phalloidin in grey. Arrows indicate overlap between ApoE- and LAMP1-positive puncta. **D.** Representative orthogonal confocal images of N2a APP_Swe_ cells treated with astrocyte-derived ApoE3 and ApoE4 for 4 h. The cells are labeled for LAMP1-positive vesicles (green), 82e1-positive Aβ/APP-βCTFs (red) and DAPI (blue) to study the localization of Aβ-containing APP processing products in N2a APP_Swe_ cells. White arrows highlights 82e1-positive puncta co-localizing with LAMP1-positive puncta. Scale bars are 5 μm (**B-D**). **E.** Schematic overview of ApoE KO, ApoE3 KI and ApoE4 KI astrocyte conditioned media collection and 4 h treatment of wild-type and AD APP/PS1 transgenic primary neurons (19 DIV). **F.** Representative confocal images of APP/PS1 neurons incubated with ApoE KO, ApoE3 or ApoE4 astrocyte conditioned media for 4 h. The intracellular intersection between internalized ApoE and endogenous human APP cleavage products was studied by labeling neurites for ApoE (green), APP metabolites Aβ/APP-βCTFs (82e1) (magenta) and MAP2 (grey). Arrows indicate ApoE and 82e1 co-localization at neurites. Scale bar is 15 μm. **G.** Higher magnification images of **Figure 4F** (indicated by a white box) of ApoE3 and ApoE4 astrocyte conditioned medium treated APP/PS1 neurons. The white arrows point at subcellular co-localization of internalized astrocyte-derived ApoE with endogenous human Aβ/APP-βCTFs. Scale bar is 6 μm. **H.** Schematic representation of synthetic 0.5 μM human Aβ_42_ and astrocyte-derived ApoE3 or ApoE4 treatment in neurons that do not endogenously produce ApoE (ApoE KO). **I.** Representative confocal microscopy images of Aβ_42_ and astrocytic ApoE-double treated primary neurons. The neurons were labeled for ApoE, APP/Aβ (6E10) and MAP2. Scale bar represents 20 μm. **J.** High magnification images of Aβ_42_ and ApoE-treated neurons of the area shown by a white box in **Figure 4I**. White arrows indicate ApoE and APP/Aβ overlap; the white circle highlights overlap of larger puncta of ApoE and APP/Aβ. Scale bar is equal to 6 μm. See also Supplementary Figure 5A-B.

### Internalized ApoE co-localizes with endogenous APP cleavage products in N2a cells and neurons

Previous research supports that intracellular Aβ is generated^21^ and in AD accumulates^18^ in the endosomal system, which internalized ApoE also accesses. To study whether internalized astrocyte-derived human ApoE and APP processing products intersect at an intracellular level, the cellular localization of human ApoE3 and ApoE4 was first examined in N2a cells overexpressing human APP containing the Swedish mutation (N2a APP_Swe_), which have abundant human APP/Aβ due to overexpression. N2a APP_Swe_ cells were treated with ApoE KO, ApoE3 or ApoE4 astrocyte conditioned medium for 4 h (**Figure 4A**) and subsequently co-labeled for human ApoE and antibody 82e1 directed at the N-terminus of human APP cleavage products Aβ and APP β-C-terminal fragment (APP-βCTF, also known as C99). Strong colocalization of ApoE and Aβ/APP-βCTF puncta were observed in N2a APP_Swe_ cells treated with astrocyte-derived human ApoE3 and ApoE4 (**Figure 4B, arrows**), but not in untransfected N2a cells or in APP_Swe_ cells treated with ApoE KO conditioned media (**Supplementary Figure 5A**), supporting the conclusion that internalized human ApoE and endogenously produced APP metabolites Aβ/APP-βCTFs intersect intracellularly.

That astrocytic ApoE was similarly internalized into LAMP1-positive late endosomes/lysosomes in untransfected N2a cells (**Figure 1D**) and N2a APP_Swe_ cells (**Figure 4C, arrows**), suggests that the overexpression of human APP with the Swedish mutation in N2a cells does not alter the subcellular localization of human ApoE. To now examine whether late endosomes/lysosomes are the subcellular compartments where human ApoE and Aβ/APP-βCTFs intersect, the cellular localization of antibody 82E1 Aβ/APP-βCTF-positive puncta was determined in N2a APP_Swe_ cells. APP-βCTF/Aβ puncta were seen to overlap with LAMP1-positive puncta (**Figure 4D, arrows**). Altogether, these data suggest that internalized astrocytic human ApoE intersects with APP metabolites Aβ/APP-βCTFs in late endosomal and/or lysosomal compartments in N2a cells.

To study whether ApoE and antibody 82e1-positive APP cleavage products Aβ/APP-βCTFs also colocalize in primary neurons, APP/PS1 transgenic neurons were treated with ApoE KO, ApoE3 or ApoE4 astrocyte media (**Figure 4E**). Remarkably, ApoE and Aβ/APP-βCTF-positive puncta were also localized together in human ApoE-treated APP/PS1 neurons (**Figure 4F-G, Supplementary Figure 5B**), indicating that added ApoE and endogenous Aβ/APP-βCTFs also intersect in primary neurons.

To further examine whether astrocytic ApoE intersects with Aβ in neurons, ApoE KO primary neurons were treated with both ApoE astrocyte conditioned media and 0.5 μM synthetic Aβ_42_ (**Figure 4H**). After 4 h, ApoE and human Aβ_42_, labeled using the human-specific Aβ/APP antibody 6E10, both localized to MAP2-positive neurites (**Figure 4I, Supplementary Figure 5C**) and co-localized along these neurites (**Figure 4J, white arrows**). Interestingly, although similar ApoE and Aβ_42_ co-localization was noted after ApoE3 and ApoE4 treatment, larger fluorescent puncta of ApoE-Aβ_42_ were observed in ApoE4-treated neurons (**Figure 4J, white circle**).

### Astrocyte medium but not ApoE genotype influences APP/Aβ levels in N2a APP_Swe_ cells

Since human ApoE derived from primary astrocytes seems to intersect intracellularly with APP cleavage products Aβ/APP-βCTFs, we next examined by Western blot whether human ApoE affects the levels of APP metabolites and whether it does so in an ApoE isoform dependent manner. Western blot showed that human APP and Aβ protein levels in N2a APP_Swe_ cells and their media were not altered by the presence of the different ApoE astrocyte media after 15 min incubation (**Supplementary Figure 9A-D**). After 4 h treatment, the time point where we observed internalized ApoE and Aβ/APP-βCTF co-localization (**Figure 4B**), APP and Aβ protein levels were also not altered in both N2a APP_Swe_ lysate and media as assessed by Western blot (**Figure 5A-C, F-H, blue panel; Supplementary Figure 6E-F**). However, we noted using Western blot that full-length APP protein levels were significantly increased while secreted APPα levels were significantly decreased in N2a cells after 8 h astrocyte media treatment, independently of the presence of human ApoE (**Figure 5A, D, F, I, orange panel; Supplementary Figure 6E-F**), since also addition of APOE KO astrocyte medium induced this effect. This suggests that the presence of astrocyte conditioned media itself, and not human ApoE, increases APP protein expression in N2a APP_Swe_ cells. ApoE3 and

**Figure 5:**
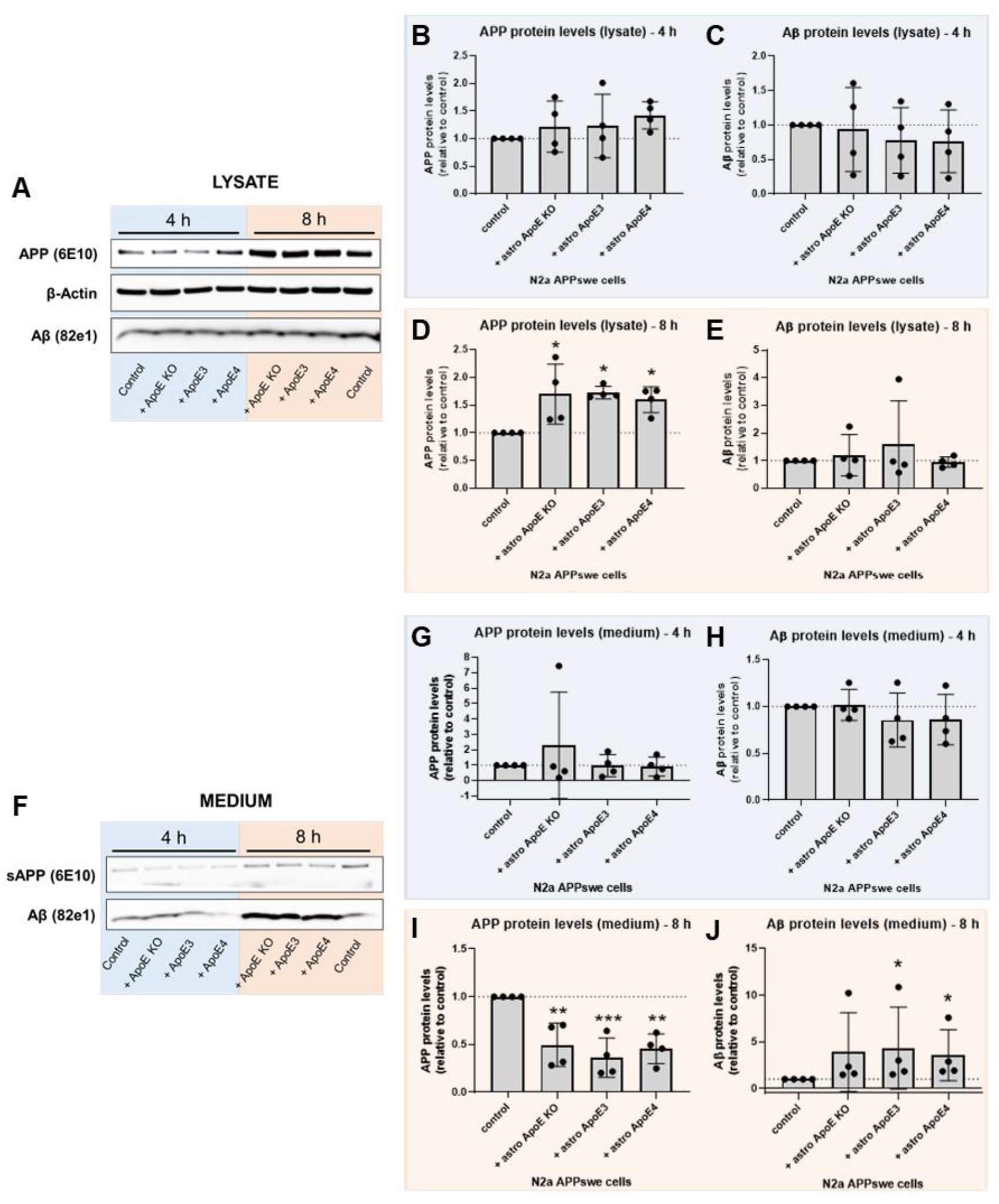
Astrocyte media but not ApoE genotype influences APP/Aβ levels in N2a APP_Swe_ cells. **A.** Representative western blot bands of lysate of N2a APP_Swe_ cells treated with ApoE KO, ApoE3 or ApoE4 astrocyte conditioned media or control (fresh N2a media)for 4 h (blue) and 8 h (orange). The western blot membranes were stained for APP, detected by 6E10 antibody, β-Actin and Aβ, detected using antibody 82e1. **B-E.** Quantification of western blots shown in **Figure 5A**. Full-length APP and Aβ protein levels were quantified in lysate of N2a APP_Swe_ cells treated with astrocyte conditioned media for 4 h (**B and C**, respectively, blue) or 8 h (**D and E,** respectively, orange). Values were normalized to β-actin and control N2a APP_Swe_ cells (APP and Aβ protein levels in these cells were set to 1). Data is shown as mean ± SD. **F.** Representative western blot membranes of sAPPα and Aβ protein levels in media from N2a APP_Swe_ cells treated with ApoE KO, ApoE3 or ApoE4 astrocyte media or control (fresh N2a media) for 4 h (blue) or 8 h (orange). **G-J.** Quantification of secreted sAPPα and Aβ protein levels in the conditions shown in **Figure 5F**. sAPPα and Aβ levels were determined in N2a APP_Swe_ cells treated with astrocyte media for 4 h (**G and H,** respectively, blue) or 8 h (**I and J,** respectively, orange). Data is shown as mean ± SD. * p-value < 0.05, ** p-value < 0.01. See also Supplementary Figure 6.

ApoE4 astrocyte conditioned media treatment did not significantly alter intracellular Aβ protein levels in N2a APP_Swe_ cells (**Figure 5A, E**), while secreted Aβ levels were significantly increased (**Figure 5F, J**); a trend for elevated secreted Aβ level was also noted at 8 h with ApoE KO astrocyte media (p-value: 0.11) (**Figure 5J**).

### ApoE4 induces increased intraneuronal Aβ_42_ levels in cultured neurons

To study whether human ApoE alters Aβ levels in primary neurons, Aβ_40_ and Aβ_42_ levels were determined in media collected from primary ApoE KO brain cultures treated with ApoE3 or ApoE4-astrocyte conditioned media for 24 h. To detect mouse Aβ in media the more sensitive technique of Mesoscale analysis was used. Interestingly, treating neurons with astrocyte conditioned media increased Aβ_42_ but not Aβ_40_ levels in neuron conditioned media in an ApoE-independent manner (**Supplementary Figure 7A-B**) and increased the Aβ_42_/Aβ_40_ ratio secreted by neurons (**Supplementary Figure 7C**).

Endogenous mouse Aβ_42_ was next investigated by immunofluorescence using Aβ_42_ antibody 12F4, as described previously^10^, in primary brain cultures obtained from ApoE KO, ApoE3-KI and ApoE4-KI mice (**Figure 6A**). Interestingly, ApoE4 neurons showed significantly higher intensity of Aβ_42_ labeling in neuronal cell bodies compared to ApoE KO neurons (**Figure 6B-C, Supplementary Figure 5D**), although no statistical difference in Aβ_42_ signal was detected between different ApoE isoforms in neurites (**Figure 6D-E, Supplementary Figure 5E**).

**Figure 6:**
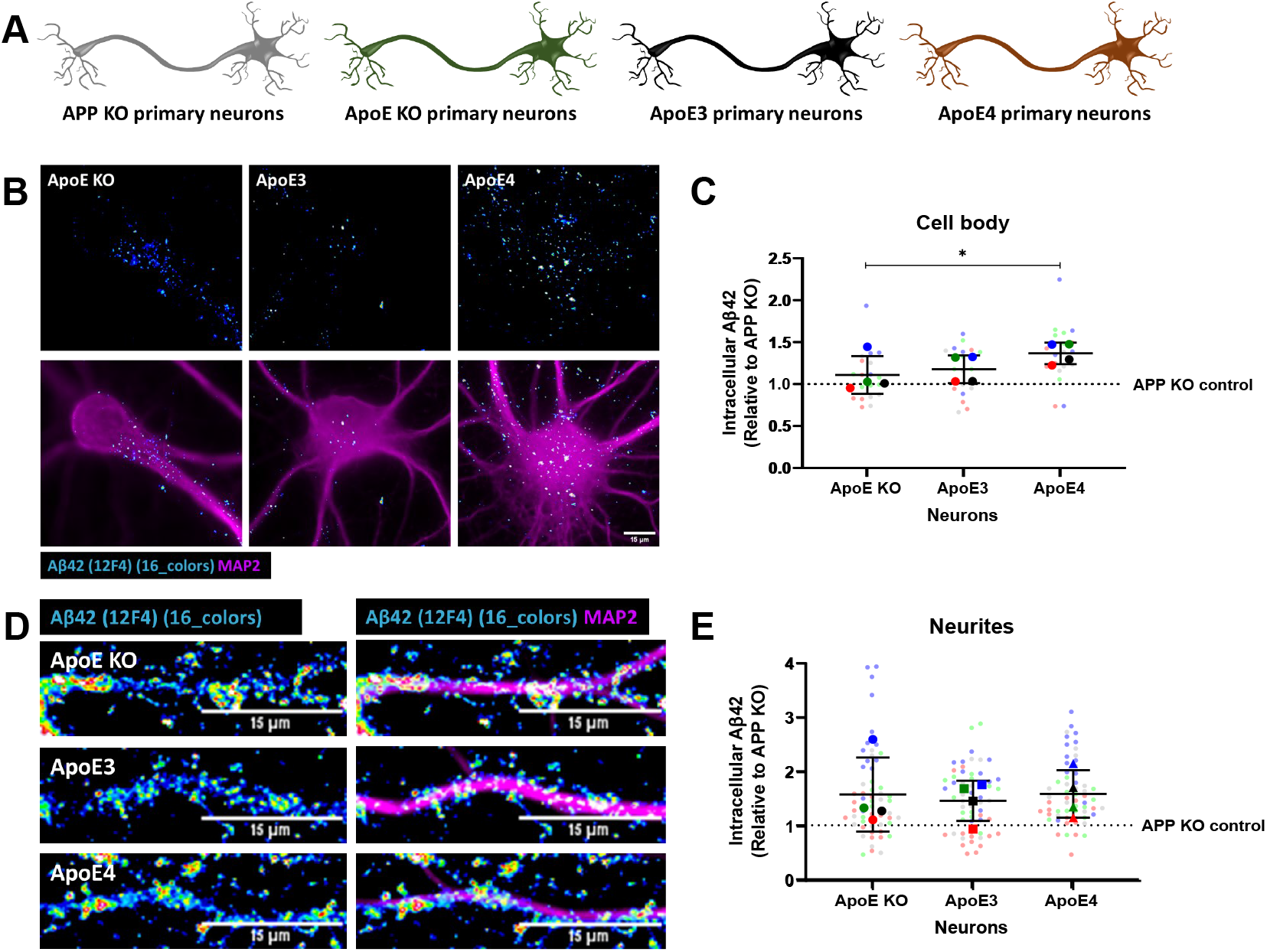
Increased endogenous intraneuronal Aβ_42_ levels in cell bodies of cultured ApoE4 neurons. **A.** Schematic visualization of the different primary neuron models used to study the effects of human ApoE isoforms on levels of intraneuronal Aβ_42_. **B.** Representative epifluorescence images of neuronal cell bodies from ApoE KO, ApoE3-KI and ApoE4-KI primary neurons (19 DIV). The neurons were labeled for endogenous mouse Aβ_42_ using antibody 12F4 (16 colors) and MAP2 (magenta). Scale bar is 15 μm. **C.** Quantification of endogenous Aβ_42_ in neuronal cell bodies as shown in **Figure 6B**. The levels of Aβ_42_ were significantly higher in ApoE4-KI neurons compared to ApoE KO neurons. **D.** Representative images of ApoE KO, ApoE3-KI and ApoE4-KI neurites labeled for Aβ_42_ by antibody 12F4 (16 colors) and MAP2 (magenta). Scale bar is equal to 15 μm. **E.** Quantification of intracellular Aβ_42_ levels, as measured by 12F4 intensity, of neurites of ApoE KO, ApoE3-KI and ApoE4-KI primary neurons. Data is shown as mean ± SD. The dashed line indicates the level of unspecific signal detected in APP KO control neurons. * p-value < 0.05. See also Supplementary Figure 5D-E.

To further examine isoform-dependent effects on intraneuronal Aβ, we treated primary human ApoE KI mouse brain cultures that endogenously express human ApoE3 or ApoE4 in both neurons and astrocytes with synthetic human Aβ_42_ for 4 h (**Figure 7A**). Of note, the number of internalized human Aβ_42_ puncta, detected using the human specific Aβ/APP antibody (6E10) were increased in ApoE4 compared to ApoE KO and ApoE3 primary brain cultures (**Figure 7B-C**, **Supplementary Figure 5F**). No difference in the area of 6E10-positive Aβ_42_ puncta was detected (**Figure 7B, D**). Strikingly, almost all the intraneuronal human Aβ puncta in both ApoE3 and ApoE4 neurons were also detected by the antibody OC against Aβ fibrils and oligomeric fibrils (**Figure 7B, lowest panel**), supporting that the internalized human Aβ_42_ is aggregated.

**Figure 7:**
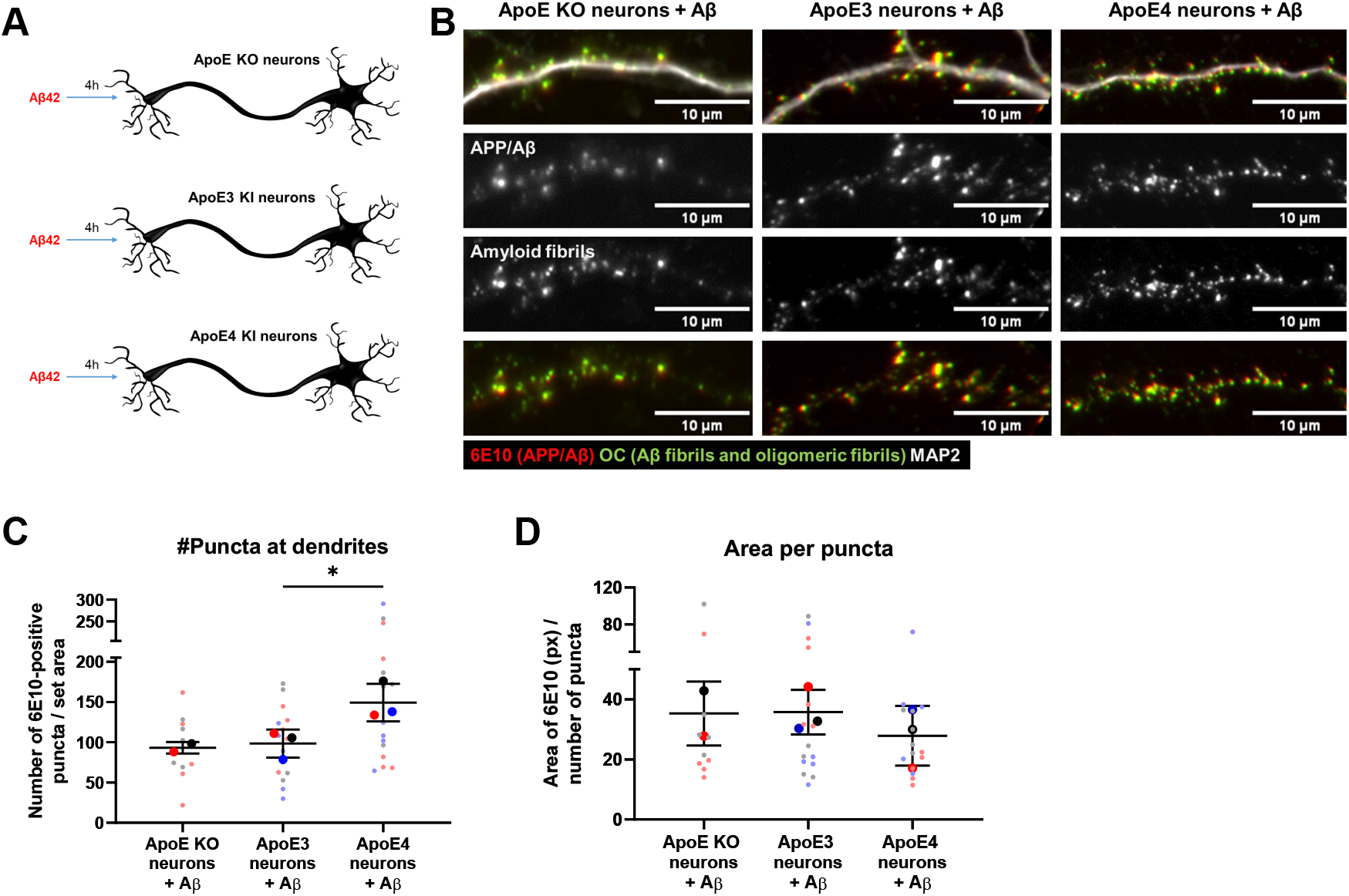
Internalized Aβ_42_ levels are increased in human ApoE4-KI primary neurons. **A.** Schematic representation of 0.5 μM synthetic Aβ_42_ treatment to different ApoE neuron cultures: ApoE KO, and human ApoE3- and ApoE4-KI neurons. **B.** Representative epifluorescence images of neurites from Aβ_42_-treated ApoE KO, ApoE3 and ApoE4 primary neurons. The neurites are labeled for human Aβ antibody 6E10 (red), fibrillar oligomer antibody OC (green) and MAP2 (grey in upper panel). Clear co-localization was observed between 6E10 and OC, suggesting that added synthetic Aβ_42_ is aggregating inside neurons. **C-D.** Quantification of the number of (**C**) and the average area size of the 6E10-positive Aβ_42_ puncta (**D**) in Aβ_42_-treated primary neurons as shown in **Figure 7B**. The number of 6E10 puncta is significantly increased in ApoE4 compared to ApoE3 neurons after Aβ_42_ treatment. The quantifications were performed while being blinded (**C-D**). Data is shown as mean ± SD. * < 0.05. px = pixels. See also Supplementary Figure 5F.

## Discussion

The prior literature suggests that ApoE4 plays a role in endosomal (dys)function^7,12,14,16,47^. However, intracellular trafficking of ApoE in normal and AD conditions remain poorly studied. We now show that ApoE is present in the endosome-autophagy-lysosome system of N2a neuroblastoma cells and in primary neurons and astrocytes, albeit in somewhat different subcellular patterns, with neurons showing endosome-autophagosome labeling particularly of neurites and less in lysosomes. We demonstrate for the first time that astrocytic human ApoE is internalized and intersects with endogenous APP cleavage products Aβ/βCTF in primary neurons and N2a APP_Swe_ cells. ApoE3 compared to ApoE4 was not seen to differentially alter endogenous APP in culture, although astrocyte media, even when devoid of ApoE, increased APP levels. However, ApoE4 increased endogenous intraneuronal mouse Aβ_42_ levels in primary neurons compared to the absence of ApoE. Additionally, ApoE4 altered intraneuronal levels of internalized exogenously added Aβ_42_ in an ApoE isoform dependent manner (ApoE4 > ApoE3). Our study highlights the importance of the endosomal system in ApoE biology and reveals an intersection between astrocyte-derived ApoE and APP processing products in neurites that we hypothesize to be of considerable importance in the pathogenesis of AD.

Due to the highly polarized shape of neurons, neuronal endosomal trafficking is quite distinct from other cell types, which might relate to our observation that ApoE clearly localized to lysosomes in N2a cells but not in neurons. In neurons, lysosomal maturation occurs during retrograde transport of endosomes-autophagosomes in neuronal processes towards the cell soma, with the most mature lysosomes being present in the soma itself^39,40^. The fact that we did not observe ApoE in lysosomes in the neuron cell soma (**Figure 2B-C**), even after BafA1 treatment (not shown), suggests that ApoE is not trafficked to and degraded by lysosomes in neurons. Previous studies^12,13,48^, including a study in primary neurons^12^, demonstrated that ApoE can be recycled, implicating endosomal recycling and re-secretion as pathways relevant to neuronal ApoE trafficking.

Our study demonstrates that astrocyte-derived human ApoE is present at APP processing sites, with ApoE and human Aβ/APP-βCTF co-localizing in neurons and neuroblastoma cells. As antibody 82E1 directed against the free N-terminus of Aβ and APP-βCTF does not differentiate between these, we cannot draw conclusions on ApoE specifically intersecting with either Aβ or APP-βCTF. The Aβ domain, however, resides in both Aβ and APP-β-CTFs and both are viewed as participating in the pathogenesis of AD. In N2a cells, ApoE and Aβ/APP-βCTF were detected in LAMP1-postive vesicles, supporting that late-endosomes and/or lysosomes are the cellular sites where ApoE intersects with Aβ/APP-βCTF. In contrast, in neurons, ApoE localized to diverse endosomal-autophagic vesicles but not to lysosomes. Due to the unique endosome-lysosome-autophagy biology in neurons^49^, additional work is required to better define the subcellular site(s) and implications of the ApoE and Aβ/APP-βCTF intersection in neurons for AD.

Kuszyczk et al.^50^ reported that inhibition of Aβ and ApoE binding affects intraneuronal Aβ levels. Thus, the intersection of Aβ and ApoE we observed in neuronal vesicles seems to affect intraneuronal Aβ. Previous papers also reported that the presence of ApoE in general is linked to increased intraneuronal Aβ compared to when ApoE is knocked out^50,51^. Huang et al.^52^ reported that human ApoE produced in HEK293T cells influences APP and Aβ secretion in an ApoE isoform-dependent manner (ApoE4 > ApoE3 > ApoE2). Here, we showed that human ApoE derived from primary astrocytes did not induce ApoE-specific effects on the secretion of APP and Aβ nor on endogenous levels of APP in N2a cells or neurons. In fact, we observed that just adding astrocyte conditioned media even devoid of ApoE to N2a cells increased APP. Our data is consistence with Huang et al, who showed that the ApoE isoform effect on Aβ production was abolished when glia were present in the culture^52^. Together, these data support the conclusion that factors in glia media markedly affect APP and Aβ levels independent of ApoE. A recent study on human iPSC-derived neurons reported an isoform-dependent astrocytic ApoE4 media increase in APP levels^53^; in contrast, in vivo studies indicate that ApoE genotype does not affect APP mRNA and protein levels in brains of humanized ApoE target replacement mice^26^ and in transgenic PDAPP mice cross-bred with ApoE target replacement mice^54^, arguing against an effect of human ApoE isoforms on APP metabolism. Wang et al.^35^ recently suggested that lipidated ApoE with cholesterol could change the localization of APP at the membrane in N2a cells towards GM1 lipid clusters, rather than changing its expression levels. As β-secretase is associated to lipid clusters, while α-secretase is linked to non-lipid-rich membrane regions, the ApoE-induced APP shift to lipid clusters in the membrane could favor Aβ production without affecting APP levels. It is also possible that the complex homeostatic control of Aβ levels might limit actual changes in levels that can be detected in cellular experiments^55^.

By adding elevated levels of Aβ_42_ to neurons, we revealed an ApoE isoform-specific effect on intraneuronal Aβ_42_, with ApoE4 inducing an increased number of intraneuronal Aβ_42_ puncta (**Figure 7**). This suggests that ApoE4 affects Aβ_42_ internalization, subsequent aggregation and/or degradaton in neurons. A similar study in N2a cells further supports this by reporting enhanced Aβ internalization when co-cultured with ApoE4-expressing cells^56^. Although the potential role of intraneuronal ApoE and its intersection with Aβ/APP-βCTF requires further study, that ApoE intersects with endogenous APP processing products as well as added Aβ, points to a role of the ApoE and Aβ interaction at different levels related to APP and Aβ biology.

In addition to the endosome-autophagosome pattern of ApoE labeling that we now describe in neurites, we previously described that added astrocyte-derived ApoE localized to synaptic terminals in primary neurons^57^. While the focus of the current paper was on internalized ApoE in neurons, ApoE in the current study was also seen in a pattern of labeling around cell bodies and neurites consistent with the synaptic localization described in our previous paper. Koffie et al.^32^ showed that ApoE co-localizes with Aβ oligomers at synapses in human brains, highlighting synaptic terminals as a potential site of ApoE and Aβ intersection. Endosomal trafficking is crucial for synaptic function^33,58^ and abnormal endosomal regulation, for example, by overexpressing Rab5, causes synaptic dysfunction^59,60^. We previously showed altered neuronal activity dependent on ApoE genotype^57^, however, it remains unclear whether and how this might relate to altered endosomal trafficking. Next to neurons, we detected ApoE internalization into primary astrocytes where it also appeared more prominent in their processes than cell soma (**Figure 3**).

In conclusion, due to the complex endosomal biology of neurons, the endosomal trafficking of ApoE is different in neurons, as internalized ApoE does not end up in lysosomes of neuronal cell bodies as it does in N2a cells and primary astrocytes. In addition, ApoE and Aβ/APP-βCTF, all key players in AD, intersect subcellularly, with isoform dependent ApoE effects. Our work further highlights the importance of the endocytic pathway in relation to the major AD players ApoE and Aβ/APP-βCTF in the early cellular phase of AD, and suggests that the endosome-lysosome-autophagy system is a potential site where ApoE and Aβ interact in AD.

## Acknowledgements

We thank Bodil Israelsson, Lund University, for her support with the animal experiments and genotyping of mice. We also thank MultiPark for the use of the confocal microscopy and Imaris software facilities. This project was funded by the European Union Horizon 2020 Research and Innovation Program SYNDEGEN (Marie Skłodowska-Curie grant agreement No. 721802), Innovation Fund Denmark (BrainStem; 4108–00008 A), the Swedish Research Council grant (No. 2019-01125), Alzheimerfonden and Konung Gustav V:s & Drottning Victorias Stiftelse.

## Author contributions

Conceptualization: S.C.K. and G.K.G.; Formal analysis: S.C.K and E.N.; Investigation: S.C.K, E.N., I.M. and L.T.G; Writing – original draft: S.C.K.; Writing – Review & Editing: S.C.K., E.N., C.G.A. and G.K.G.; Visualization; S.C.K. and E.N.; Supervision: C.G.A. and G.K.G.; Funding Acquisition: G.K.G.

## STAR methods

### RESOURCE AVAILABILITY

#### Lead contact

Further information about the resources and reagents used in this study should be directed to the lead contract, prof. Dr. Gunnar K. Gouras.

#### Material availability

In this study, now new reagents were generated.

#### Data and code availability

All data reported in this paper will be shared by the lead contact upon request. This paper does not report original code. Any additional information required to reanalyze the data reported in this paper is available from the lead contact upon request.

### EXPERIMENTAL MODEL AND SUBJECT DETAILS

#### Animals

In this study, the following mouse models were used: ApoE KO (B6.129P2-Apoe<tm1Unc>/J, the Jackson Laboratory), humanized ApoE3-KI (B6.Cg-Apoeem2(APOE*)Adiuj/J, the Jackson Laboratory), humanized ApoE4-KI (B6(SJL)-Apoetm1.1(APOE*4)Adiuj/J, the Jackson Laboratory), and AD transgenic APP/PS1 B6.Cg-Tg mice (APPswe, PSEN1dE9)85Dbo/Mmjax. All animal experiments described in this study were approved by the Ethical Committee for animal research at Lund University, Sweden (permit number: M5983-19).

#### Primary mouse mixed brain cultures

Primary mouse brain cultures were obtained from cortical and hippocampal brain tissue from ApoE KO, ApoE3-KI, ApoE4-KI, wild-type and APP/PS1 embryos (E15-E17). Embryonic brain tissue was dissected and dissociated into single cells as previously described by Takahashi et al.^61^. In short, cortical and hippocampal embryonic brain tissue was manually dissociated using 0.25% trypsin (Thermo Fisher Scientific, 15090046) and seeded onto poly-D-lysine coated coverslips or plates. Directly after seeding, the cells were cultured in Dulbecco’s modified Eagle medium (DMEM) containing 10% fetal bovine serum (FBS) (Gibco, 10082147) and 1% antibiotics penicillin/streptomycin (p/s) (Thermo Fisher Scientific, SV30010). After 3-5 hours, the 10% FBS medium was replaced by complete Neurobasal medium containing B27 supplement (Gibco, 17504044), 1% p/s and 1.4 mM L-glutamine (Gibco, 25030081) and cultured for 19 days *in* vitro (DIV) until further use. Both male and female embryos were used to generate primary brain cultures, although we did not specifically identify the sex of embryos; our blinding to gender also could reduce potential bias in subsequent analyses. Our experience indicates that offspring of our crosses as expected generate about equal number of male and female offspring.

#### Primary mouse astrocytes

Primary mouse astrocyte cultures were obtained from ApoE KO, ApoE3-KI and ApoE4-KI mouse pups (P1-P3) as previously described by Konings et al.^57^. In short, cortical and hippocampal brain tissue was obtained from mouse pup brains and manually dissociated after 0.25% trypsin incubation using plastic Pasteur pipets. After dissection and dissociation of the tissue, cells were seeded on T75 plates coated with poly-D-lysine. Primary astrocytes were cultured in AstroMACS medium (Miltenyi Biotec, 130-117-031) with 0.5 mM L-glutamine and the medium was replaced every 2-3 days. Analogous to primary brain cultures, also for our generation of primary astrocytes we did not separate cells based on gender, but used all available pups which should lead to analogous number of male and female cells.

#### Culturing neuroblastoma N2a cells

Mouse neuro-2a (N2a) cells (ATCC^®^ CCL-131™) without transfection or N2a cells stably transfected with human APP carrying the Swedish mutation (N2a APP_Swe_)^62^ were cultured in DMEM and Opti-MEM (Gibco, 31985062) medium (ratio 1:1) containing 10% FBS and 1% p/s at 37°C and 5% CO_2_. N2a cells transfected with human APP_Swe_ were selected using 50 mg/mL Geneticin (Gibco, 10131027) in their medium. N2a and N2a APP_Swe_ cells were seeded on poly-D-lysine coated glass coverslips one day prior to recombinant or astrocytic ApoE treatment.

### METHOD DETAILS

#### Recombinant ApoE treatment

Recombinant ApoE3 and ApoE4 proteins (Sigma-Aldrich, SRP4696 and A3234, respectively) were reconstituted in 0.1% bovine serum albumin (BSA) in milli-Q water to a stock concentration of 0.1 mg/ml. All experiments used a final ApoE concentration of 2.5 μg/ml.

#### Astrocyte conditioned media treatment

Once primary astrocyte cultures reached 80% confluence, ApoE KO, ApoE3 and ApoE4 astrocyte conditioned media were collected. For the medium collection, astrocytes were first shortly washed with phosphate buffered saline (PBS) and cultured in complete Neurobasal medium. After 48 hours, astrocyte conditioned media were collected on ice, centrifuged at 10,000 rpm at 4°C for 10 min and supernatant stored at −80°C in small aliquots to avoid freeze-thaw cycles.

Astrocyte conditioned media from ApoE KO, ApoE3 and ApoE4 astrocyte cultures were used to treat primary neuron cultures or mouse neuroblastoma neuro-2a (N2a) cells. Half of the original culture medium from primary neurons and N2a cells was replaced by an equal volume of conditioned astrocyte medium, resulting in half astrocyte conditioned and half Neurobasal/N2a medium.

#### Lysosome inhibition experiments

To block lysosomal function in neurons, lysosomal acidification inhibitor Bafilomycin A1 (Sigma) was used. Bafilomycin A1 was reconstituted in DMSO and further diluted to a final concentration of 10 nM to treat primary neurons. Primary neurons were treated 1 h prior to ApoE addition for a total duration of 5 h before fixation.

#### Synthetic Aβ_42_ treatment

Synthetic Aβ_42_ peptides (Tocris) were prepared as previously described by Klementieva and colleagues^63^. In short, synthetic Aβ_1-42_ was reconstituted in DMSO to a stock volume of 250 μM. The peptides were further prepared by sonicating for 10 min followed by centrifuging at 12,000 rpm for 15 min. An end concentration of 0.5 μM Aβ_42_ was used to treat primary neurons.

#### Western blot

Western blot analysis was performed on astrocyte-conditioned media collected from ApoE KO, ApoE3-KI and ApoE4-KI primary astrocytes to confirm the presence of human ApoE in ApoE3 and ApoE4 astrocyte conditioned media. Medium was directly collected from astrocytes and centrifuged at 10,000 x g for 10 min at 4°C. The supernatant was collected and directly used or stored at −80°C. Medium samples were prepared for loading by mixing with NuPage reducing agent (Invitrogen, NP004), and Novex NuPage LDS sample buffer (Invitrogen, NP007). Subsequently, the samples were heated at 70°C for 10 min and centrifuged prior to loading to a NuPAGE 4-12% Bis-Tris gel (Invitrogen, NP0321BOX).

Western blot was also performed on N2a APP_Swe_ cells treated with ApoE astrocyte-conditioned media to determine the APP and Aβ protein levels after human ApoE treatment. 4 h after astrocyte media treatment, media of N2a cells was first collected and N2a cells were washed in PBS, gently scraped, collected in a 1.5 ml tube and centrifuged at 10,000 x g for 2 min. The supernatant was removed and the pellet was either snap frozen at −80°C in liquid nitrogen until further use or directly used for western blot. To detect Aβ using western blot, lysate samples were lysed in PBS containing 6% SDS and 1% β-mercapto-ethanol, followed by sonication, heating at 95°C for 6 min and centrifugation at 12,000 rpm for 10 min. To detect Aβ, the N2a lysate samples were mixed with Novex™ Tricine SDS Sample Buffer (Invitrogen, LC1676), boiled at 95°C for 5 min, shortly centrifuged and loaded onto a Novex™ 10 to 20% Tricine gel (EC6625BOX).

SDS PAGE gel electrophoresis, the proteins were transferred to a PDVF membrane using iBlot™ 2. For APP and Aβ detection, the membranes were subsequently washed in PBS and cut into two (cut was made around 16 kDa). The membrane part containing <16 kDa proteins was boiled for 5 min in PBS to improve the detection of Aβ. The membrane part containing >16 kDa proteins was stained for total proteins (Azure, AC2225) accordingly to manufacturer’s instructions and images were acquired using a Sapphire Biomolecular imager (Azure Biosystems). After total protein quantification, the top part of the membrane (>16 kDa) was further cut just below 78 kDa to obtain a total of three membrane parts (**Supplementary Figure 6E-F**). All membranes were blocked in PBS-T containing 5% skim milk powder. The membranes were incubated overnight with primary antibodies at 4°C, followed by secondary HRP-conjugated antibodies for 1 hour at room temperature. All washes were done in PBS-T. The membrane were developed using ECL Substrate and visualized using a Sapphire Biomolecular imager.

Quantification of the bands was performed using Image Lab 6.1. All bands were normalized to total protein and β-actin. For protein quantification in media, the bands were corrected for both total proteins in media as well as β-actin and total protein from its corresponding lysate to correct for both protein concentrations in medium as well as differences in cell number. All bands are presented are normalized to the control, where the control condition is untreated N2A APP_Swe_ cells that were used to establish a baseline for the APP metabolism.

#### Mesoscale analysis

Secreted Aβ_40_ and Aβ_42_ levels were measured using Meso Scale Discovery (MSD). Mouse primary brain cultures from wild-type or APP/PS1 incubated for 24 h with control (fresh neurobasal media), ApoE3 KI, or ApoE4 KI astrocyte conditioned media were collected. Aβ levels in media were assessed with the 4G8 kit (#K15199E) following the manufacturer’s protocol. Plates were read using a QuickPlex Q120 reader (Meso Scale Diagnostics). Total protein in cell lysates was determined by BCA assay and used to normalize the Aβ levels to the concentration of total proteins.

#### Immunofluorescence

N2a cells and primary neurons (19 DIV) grown on glass coverslips were fixed in 4% paraformaldehyde (PFA), 4% sucrose in PBS for 15 minutes at room temperature. The cells were incubated in PBS containing 1% BSA, 0.1% Saponin (Sigma-Aldrich, 84510) and 2% normal goat serum (Jackson ImmunoResearch, 005-000-121) for 1 h at room temperature to permeabilize the cells and block unspecific signal. Afterwards, the coverslips containing N2a cells or primary neurons were incubated with primary antibodies, diluted in 2% normal goat serum in PBS, overnight at 4°C, followed by incubation with secondary fluorescently-labeled antibodies for 1 h at room temperature in the dark. The coverslips were mounted on glass slices with ProLong™ Diamond Antifade Mountant (Invitrogen, P36961). All washes were performed in PBS.

#### Epifluorescence microscopy

Microscopy was performed on coverslips containing fluorescently labeled N2a cells or primary neurons. In our experiments, two different epifluorescence microscopes have been used: 1. Olympus IX70 microscope equipped with a 405, 488, 568 and 647 nm channel, X-Cite^®^ 120Q excitation light source (Excelitas Technologies), a C11440 ORCA-Flash4-oIT digital camera, and 40x NA 1.3 and 60x NA 1.4 oil immersion objectives. An additional 1.5 magnification of the 60x objective was used when taking the images. 2. Nikon Eclipse 80i upright microscope (RRID:SCR_015572) equipped with a 10x (Nikon plan apo, NA 0.45), 20x (Nikon plan apo, NA 0.75), 40x (Nikon plan apo, NA 1.0, oil immersion), and 60x objectives (Nikon apo VC, NA 1.40, oil immersion). The Nikon microscope was connected to a computer containing Nikon Instruments Software-Elements Advance Research (NIS-Elements AR) version 3.2. The focus of the images was set based on the signal in the ApoE channel.

#### Confocal microscopy

To obtain image stacks a laser scanning confocal microscope Leica TCS SP8 (Leica Microsystems, RRID: SCR_018169) in combination with Leica Application Suite X (LAS X) software version 3.4.7.23225 (Leica Microsystems, RRID: SCR_013673) was used. A Z-step size of 0.5 μm was used for imaging stacks. Orthogonal images were obtained by using Bitplane Imaris viewer version 9.5.1 (Oxford Instruments, RRID: SCR_007370).

#### Image analysis

The co-localization of added recombinant ApoE3 and ApoE4 with sub-cellular markers Rab7, LAMP1, GM130 and TGN38 in N2a cells (**Figure 1, Supplementary Figure 2**) was analysed by determining the percentage of ApoE pixels that co-localized with each endosomal marker using Fiji ImageJ. The images were pre-processed using background subtraction and brightness processing to remove unspecific signal.

For Rab7, which was detected as small puncta with high background in the cell, the images were also sharpened before background subtraction. Pre-processing settings of ApoE was set based on control-treated N2a cells to get rid of endogenous mouse ApoE. After pre-processing of ApoE and the subcellular markers in the images, the images were thresholded and a selection was created for ApoE. Subsequently the pixel co-localization was measured in Fiji ImageJ. N2a cells containing 5 to 20 ApoE puncta per cell were included in the pixel co-localization analyses.

To study whether ApoE in neurons is degraded by lysosomes, the lysosomal activity was inhibited using bafilomycin A1 (**Figure 2J-K**). Images taken from bafilomycin- and ApoE-treated neurons were analyzed using Fiji ImageJ. First, 20 different, non-overlapping regions of interest were selected along MAP2-positive neurites for each analyzed neuron. The area of each individual region of interest was kept constant. Region of interests of neurites were selected in such a way that all intracellular ApoE will be detected, meaning that synaptic ApoE puncta proximal to but not overlapping with MAP2 labeling was also included in the analysis. Within these regions of interest, the co-localization between ApoE and LAMP1 was analyzed in the same way as described for N2a cells (previous section).

The number of MAP2-positive neurons and GFAP-positive astrocytes in our primary cultures (**Figure 3C**) was determined by manual counting using the Cell Counter plugin in ImageJ (https://imagej.nih.gov/ij/plugins/cell-counter.html). DAPI nuclei were used to determine the total number of cells in the culture.

Intraneuronal Aβ_42_ levels (**Figure 6C, E**) were calculated based on the intensity of Aβ_42_-specific antibody 12F4 as previously described by Ubelman et al.^10^. Region of interests were set for cell bodies and neurites based on MAP2 labeling using the polygon tool in ICY (www.icy.bioimageanalysis.org). The mean intensity of 12F4 labeling was measured for the selected cell bodies and neurites and were corrected for the mean intensity of background signal. For one embryo, five independent neurons were analyzed. In each analyzed neuron, the three most prominent neurites were included in the analysis. All mean intensities measured in this study were normalized to the mean intensity measured in APP KO cell bodies and neurites.

The levels of internalized synthetic Aβ_42_ in primary neurons were analyzed based on 6E10 antibody labeling (**Figure 7C-D**). The 6E10 antibody is known to exclusively label human Aβ and not endogenous mouse Aβ. For quantification of the number of 6E10 puncta and the area of the 6E10 puncta, images obtained by epifluorescence microscopy were pre-processed and thresholded using Fiji ImageJ. 10 region of interest of neurites per cell were selected using the polygon tool in ImageJ based on MAP2 labeling. Both number and size of the puncta were analyzed using particle analyzer. The puncta numbers were corrected for the total area analyzed to have all values corrected to a set area of 1000 μm^2^.

### QUANTIFICATION AND STATISTICAL ANALYSIS

All statistical analyses were performed in Graphpad Prism 8.4.1. Prior to statistical testing, it was assessed whether the data was normally (Gaussian) distributed based on normality tests Shapiro-Wilk tests and Kolmogorov-Smirnov tests, and QQ plots. In case of a normal distribution of the data, unpaired student’s t-test were performed to compare ApoE3 to ApoE4 (**Figure 1E, G**) and one-way ANOVA to compare more than two groups (**Figure 5B-D, 5H-I, 7C-D, and Supplementary Figure 7B-C**). When the data was not normally distributed, non-parametric Mann-Whitney tests were performed to test statistical difference between ApoE3 and ApoE4 (**Figure 2J-K and Supplementary Figure 2B, D**). When more than two groups were compared, Kruskal-Wallis tests was used to analyze non-normalized data (**Figure 5E, G, J, 6C, E and Supplementary Figure 7A**). All data displayed in graphs were shown as mean ± SD. Big data points in graphs reflect number of embryos (N), smaller data points reflect single neurites/cells analyzed (n).

