## Supplemental figures for "Apolipoprotein E intersects with amyloid-β within neurons"

**Supplementary Figures**

**Supplementary Figure 1**

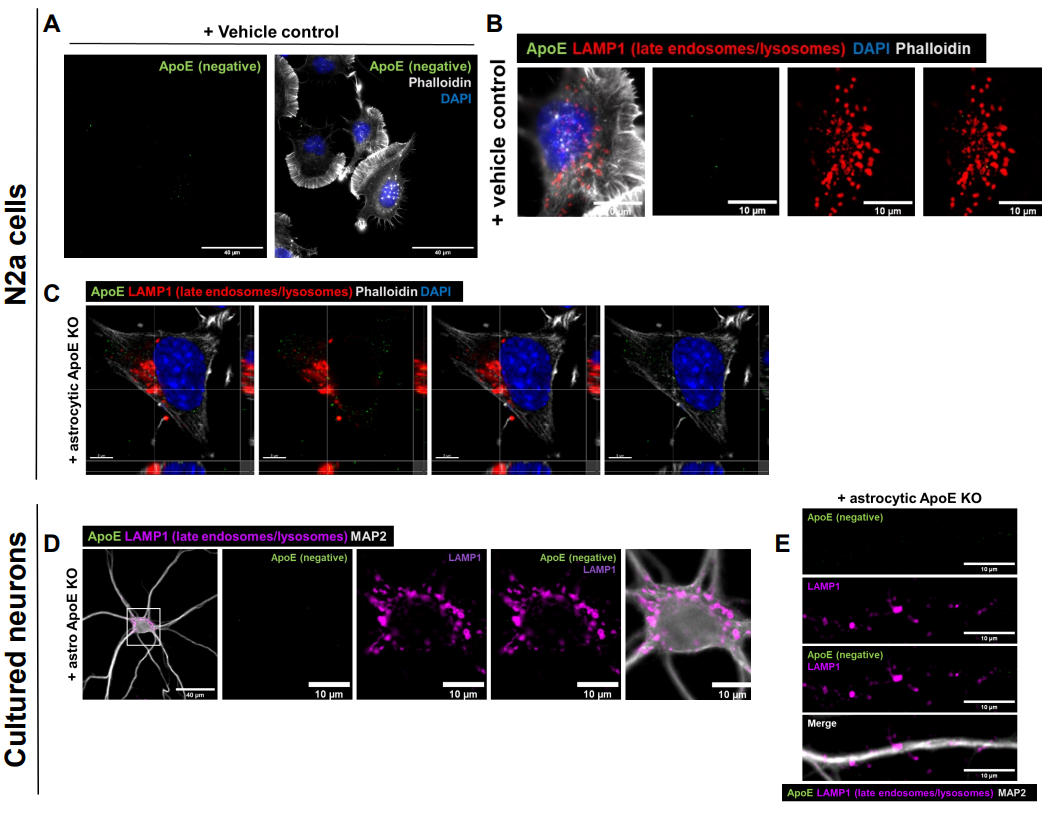

**Supplementary Figure 1: Added human ApoE cannot be detected in N2a cells or ApoE KO primary neurons treated with vehicle control or ApoE KO ACM for 4 h.** **A.** Representative epifluorescence image of N2a cells treated with a vehicle control for 4 h. No ApoE was detected in the N2a cells labeled for ApoE (green), DAPI (blue) and phalloidin (grey). Scale bar is 40 µm. See also Figure 1B. **B.** Representative images from epifluorescence microscopy of N2a cells treated with vehicle control. The N2a cells were labeled with antibodies for ApoE (green), LAMP1 (red) and stained for DAPI (blue) and phalloidin (grey). Scale bar is 10 µm. See also Figure 1D. **C.** Representative confocal images of ApoE KO ACM-treated N2a cells. The N2a cells were labeled for ApoE (green), LAMP1 (red), DAPI (blue) and phalloidin (grey). Scale bar is 5 µm. See also Figure 1K. **D.** Representative fluorescence images of ApoE KO ACM-treated primary neurons (19 DIV). The neurons were labeled for ApoE (green), neuronal marker MAP2 (grey) and LAMP1 (magenta). Scale bars are 40 µm (left panel) and 10 µm (middle and right panels). See also Figure 2B. **E.** Representative epifluorescence images of neurites from ApoE KO ACM-treated primary neurons. The neurons were labeled for ApoE (green), MAP2 (grey) and LAMP1 (magenta). Scale bars are 10 µm. See also Figure 2D.

**Supplementary Figure 2**

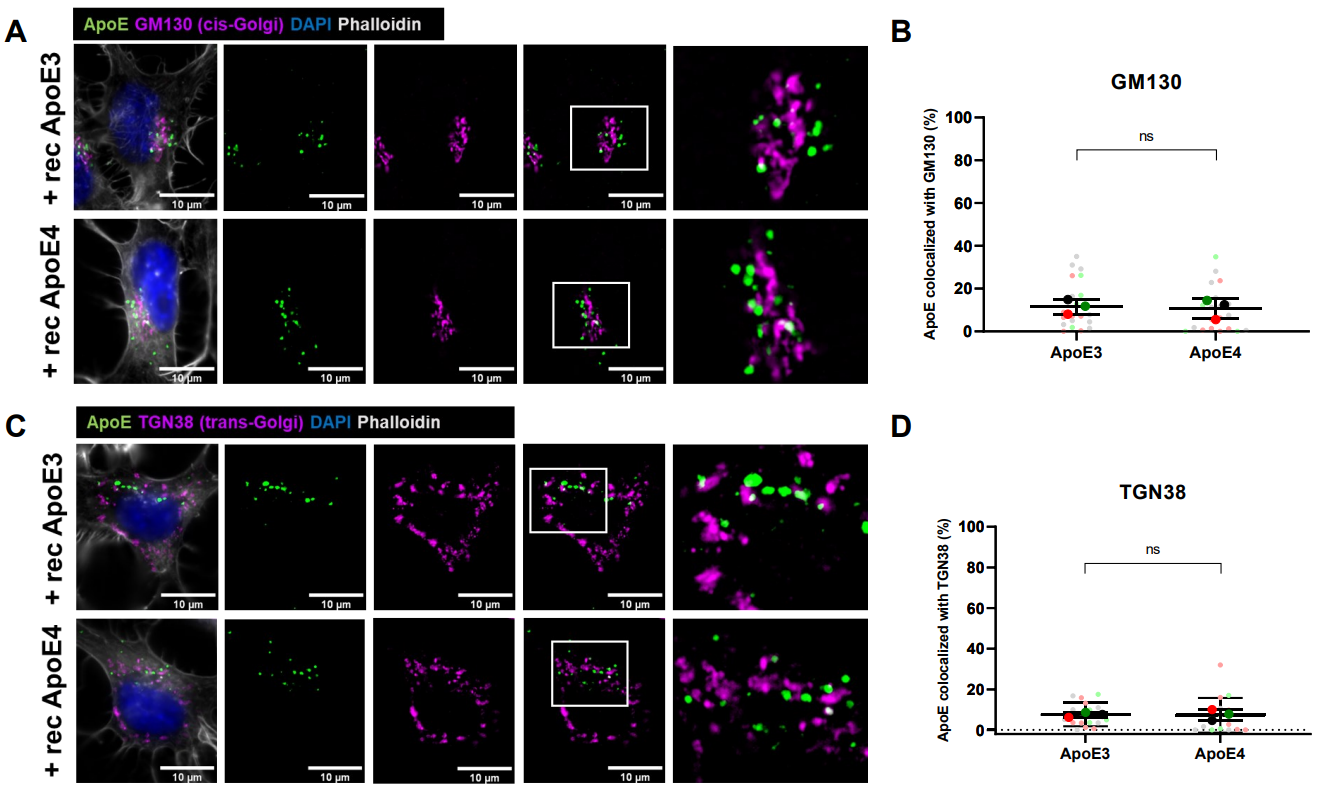

**Supplementary Figure 2: Added human recombinant ApoE is not localized at the cis- or trans-Golgi apparatus in N2a cells. A and C.** Representative epifluorescence microscopy of N2a cells treated with recombinant Apo3 or ApoE4 for 4 h. The cells were labeled for ApoE (green), cis-Golgi marker GM130 **(A)** or trans-Golgi marker TGN38 **(C)** (magenta), DAPI (blue) and phalloidin (grey). The right panels are showing a higher magnification image of the area indicated by the white box. **B and D.** Quantification of GM130 **(B)** and TGN38 **(D)** co-localization with ApoE in N2a cells treated with different recombinant ApoE isoforms for 4 h. The graphs correspond to the representative images shown in **Supplementary Figure 2A** and **Supplementary Figure 2C**, respectively. ns = not significant, rec ApoE = recombinant ApoE.

**Supplementary Figure 3**

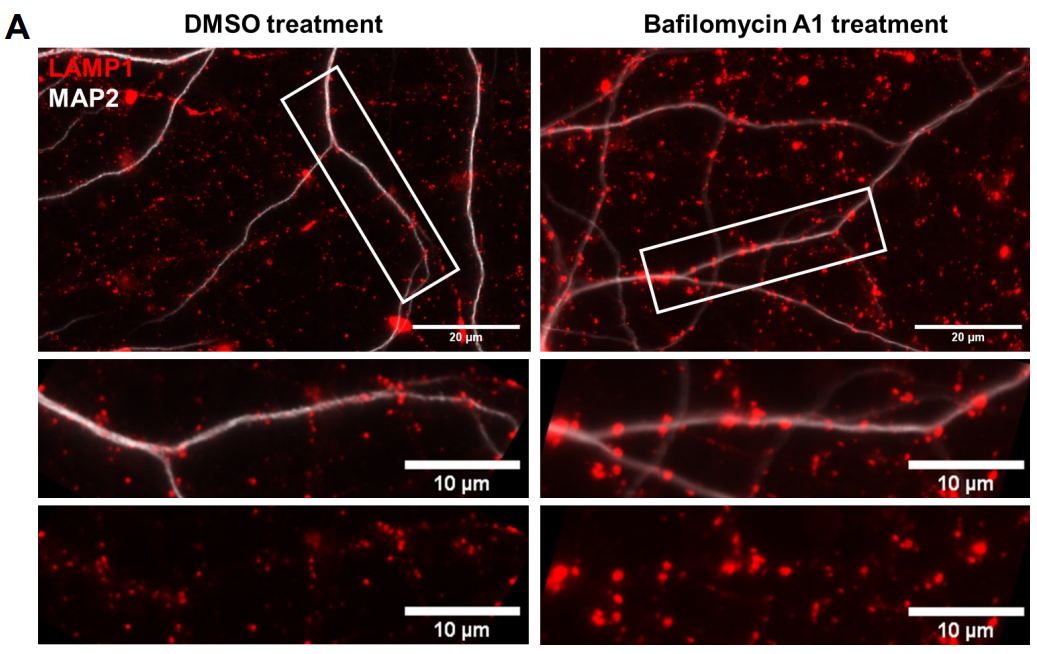

**Supplementary Figure 3: Bafilomycin A1 treatment alters LAMP1-labeled puncta in primary neurons.
A.** ApoE KO primary neurons were treated with 10 nM bafilomycin for 5 h to inhibit lysosomal activity, prior to ApoE KO astrocyte media treatment for 4 h. After the bafilomycin treatment, LAMP1 puncta was observed to be bigger and brighter compared to DMSO treated control neurons. LAMP1 was labeling late endosomes/lysosomes in red, MAP2 was labeling dendrites in grey. Scale bar is 20 µm in the top panels and 10 µm in the lower panels. The same settings were used as in **Figure 2H-I**.

**Supplementary Figure 4**

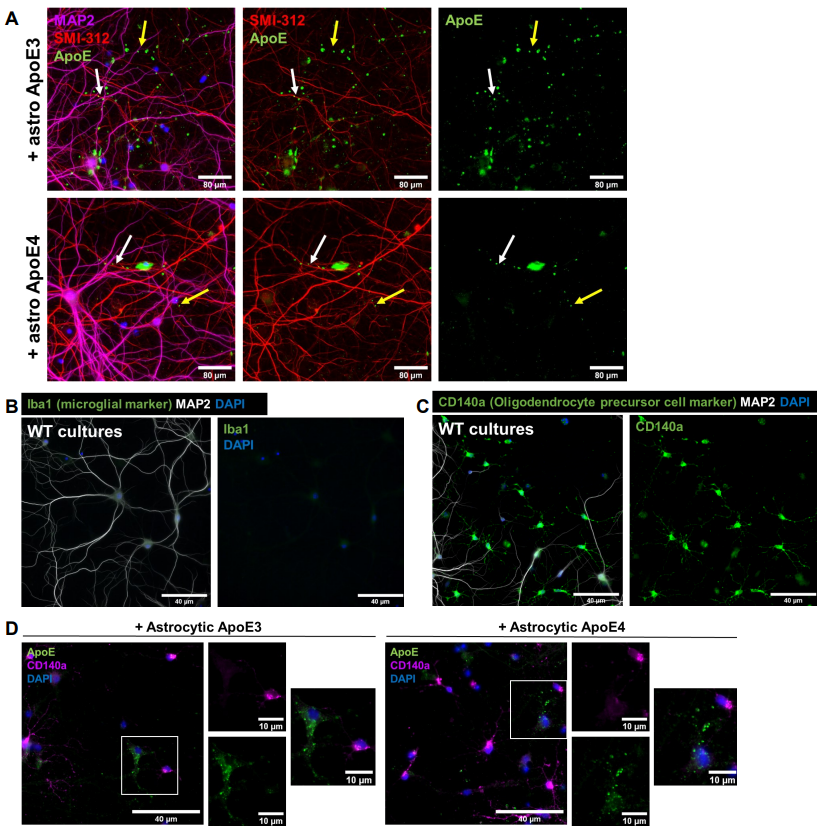

**Supplementary Figure 4: Internalized human ApoE is not purely following an axonal pattern and is not present in oligodendrocyte precursor cells.** **A.** Representative fluorescence images of primary neurons treated with astrocyte conditioned media for 4 h showing that ApoE (green) overlaps with axonal marker SMI-312 (red) to some extent (white arrow), but not always (yellow arrow). The neurons were also labeled for dendrites using MAP2. **B.** Representative fluorescence image showing that wild-type primary cultures used in this study do not contain Iba1-positive microglia. Primary mixed cultures were obtained from embryonic mouse brains. **C.** Representative microscopy images of wild-type primary cultures showing that our cultures also contain CD140a-positive OPCs (green). In addition to cellular markers, wild-type primary neurons were also labeled for neuronal/dendritic marker MAP2 and nuclear stain DAPI (**B-C**). **D.** Representative epifluorescence images of ApoE KO neurons treated with ApoE3 and ApoE4 astrocyte conditioned media for 4 h. These images show that ApoE3 and ApoE4 (green) were not present in OPCs (magenta) in primary culture. Scale bars are equal to 80 µm (**A**), 40 µm (**B, C and D, bigger panels**) and 10 µm (**D, smaller panels**). Astro = astrocytic, WT = wild-type.

**Supplementary Figure 5**

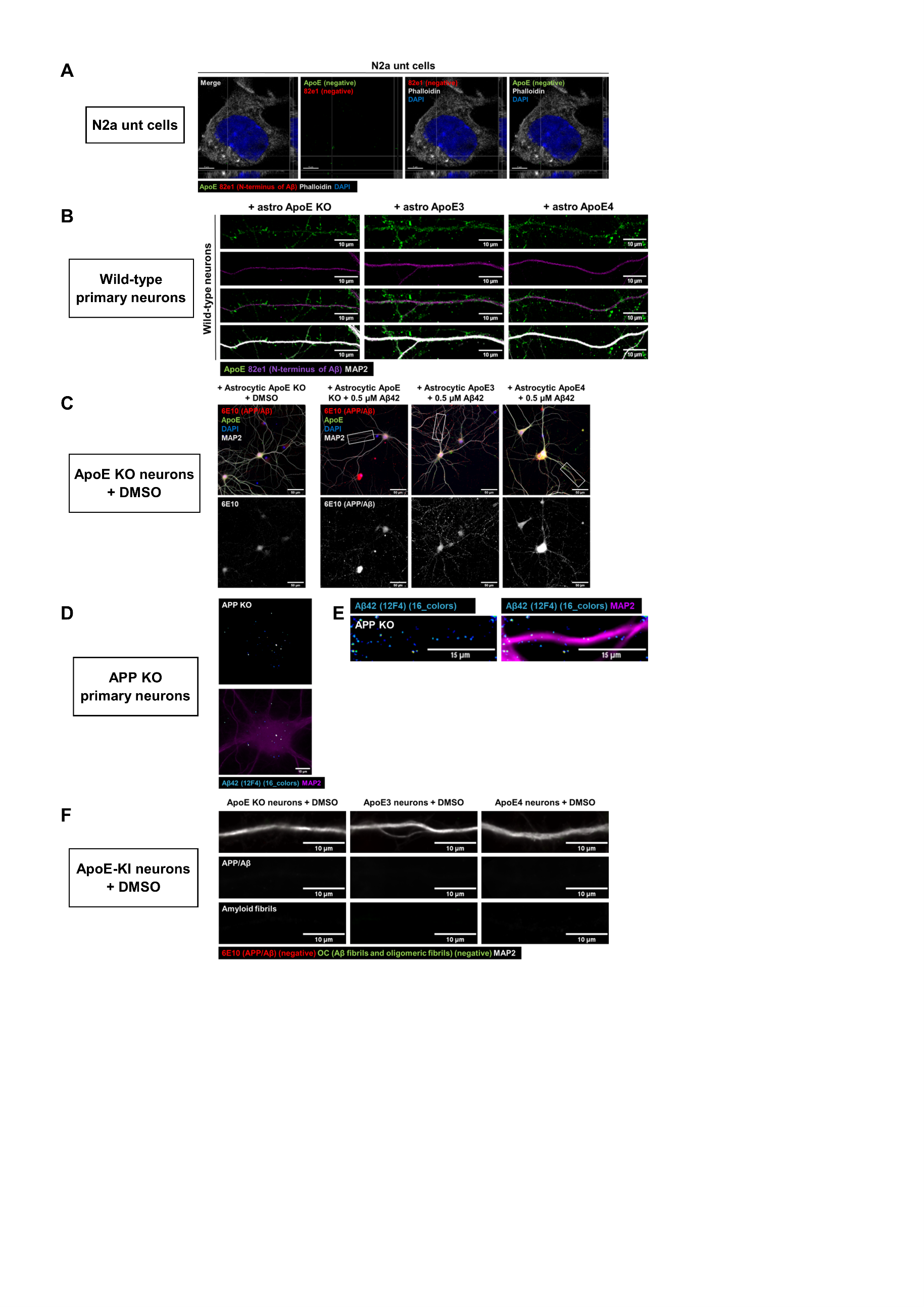

**Supplementary Figure 5: N2a untransfected cells, wild-type and APP KO neurons and cultured neurons treated with DMSO (synthetic Aβ control) show minimal human Aβ labeling.** **A.** Representative confocal images of non-treated and untransfected N2a cells. These cells were labeled for phalloidin (grey), DAPI (blue), ApoE (green) and 82e1 (red). Neither ApoE nor 82e1 labeling was detected in these cultures. Scale bar is 5 µm. See also Figure 4B. **B.** Representative epifluorescence images of wild-type neurons treated with astrocyte media. No clear labeling for the 82e1 antibody was detected in these neurons. Scale bar is 15 µm. See also Figure 4F-G. **C.** Representative confocal microscopy images of synthetic Aβ (labeled by 6E10 antibody) in ApoE KO primary neurons treated with ApoE KO astrocyte conditioned media and DMSO (left) in comparison to ApoE KO neurons treated with astrocytic human ApoE and 0.5 µM synthetic Aβ. DMSO treated neurons show lower 6E10-positive labeling. Scale bar is equal to 50 µm. See also Figure 4I-J. **D-E.** Representative images of neuronal cell bodies (**D**) and neurites (**E**) showing low/unspecific signal of Aβ_42_ antibody 12F4. The neurons were labeled for 12F4 (16 colors) and MAP2 (magenta). Scale bar are 15 µm (**D-E**). See also Figure 6B, D. **F.** Representative images of DMSO-treated ApoE KO, ApoE3 and ApoE4 primary neurons showing that these cells were 6E10- (red) and OC- (green) negative. The same settings were used as for Figure 7B. Dendrites were labeled with MAP2 (grey). Scale bar is 10 µm.

**Supplementary Figure 6**

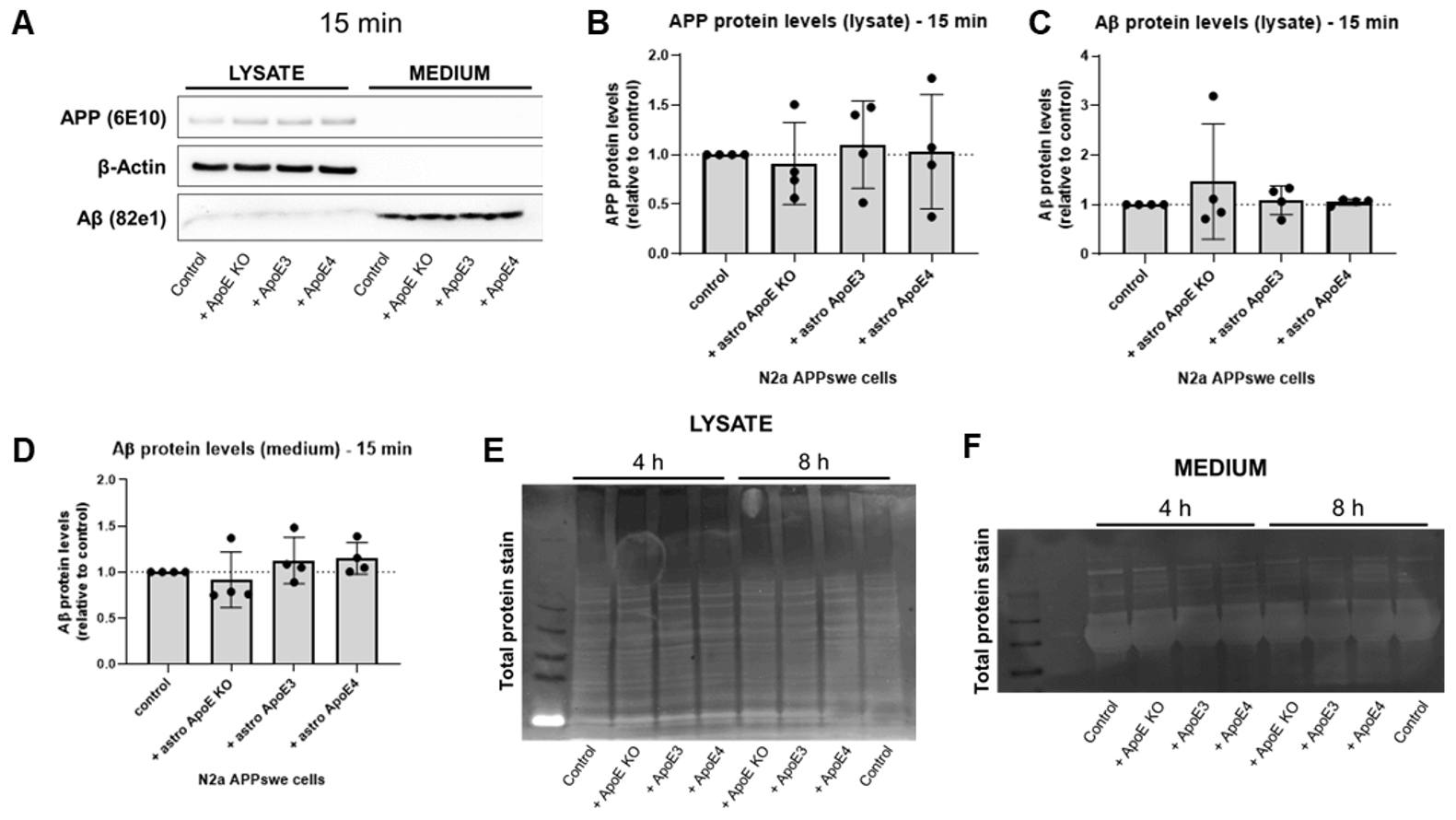

**Supplementary Figure 6: Astrocyte-derived human ApoE does not alter APP or Aβ levels in N2a APP_Swe_ cells after 15 min treatment. A.** Representative western blot membrane of APP, β-actin and Aβ protein detection in lysate and media from N2a APP_Swe_ cells treated with ApoE KO, ApoE3 or ApoE4 astrocyte conditioned media for 15 min. Control treatment reflects just media change to regular N2a media. **B-D.** Quantification of APP (**B**) and Aβ (**C**) protein levels in N2a lysate, and secreted Aβ protein levels in N2a media (**D**) after 15 min astrocyte conditioned media treatment. Data is shown as mean ± SD. **E-F.** Representative images of total protein stain of the membranes showed in Figure 5A **(E)** and 5F **(F)**.

**Supplementary Figure 7**

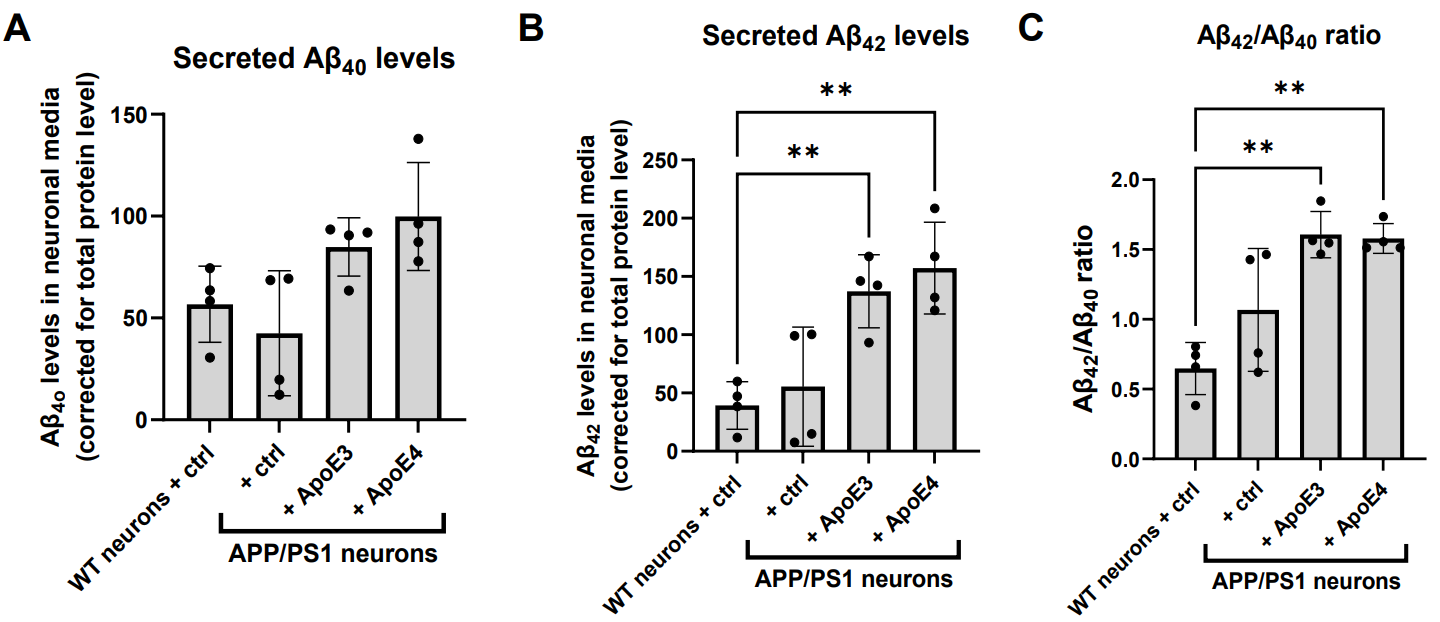

**Supplementary Figure 7: Secreted Aβ_42_ levels are increased in media from APP/PS1 primary neurons treated with astrocytic ApoE3 and ApoE4. A-C.** Graphs showing the level of secreted Aβ_40_ (**A**), Aβ_42_ (**B**), and Aβ_42_/Aβ_40_ ratio (**C**) in wild-type and APP/PS1 primary neurons treated with control (fresh neurobasal media) or ApoE3 or ApoE4 astrocyte conditioned media for 24 h. The Aβ levels were measured using mesoscale analysis and were normalized to total protein levels as measured by BCA assay. Data are shown as mean ± SD. One data points reflects one embryo. ** p-value < 0.01.
